# Circular RNA Vaccines against SARS-CoV-2 and Emerging Variants

**DOI:** 10.1101/2021.03.16.435594

**Authors:** Liang Qu, Zongyi Yi, Yong Shen, Liangru Lin, Feng Chen, Yiyuan Xu, Zeguang Wu, Huixian Tang, Xiaoxue Zhang, Feng Tian, Chunhui Wang, Xia Xiao, Xiaojing Dong, Li Guo, Shuaiyao Lu, Chengyun Yang, Cong Tang, Yun Yang, Wenhai Yu, Junbin Wang, Yanan Zhou, Qing Huang, Ayijiang Yisimayi, Yunlong Cao, Youchun Wang, Zhuo Zhou, Xiaozhong Peng, Jianwei Wang, Xiaoliang Sunney Xie, Wensheng Wei

## Abstract

Severe acute respiratory syndrome coronavirus 2 (SARS-CoV-2) and its emerging variants of concern (VOC), such as Delta (B.1.617.2) and Omicron (B.1.1.529), has continued to drive the worldwide pandemic. Therefore, there is a high demand for vaccines with enhanced efficacy, high thermostability, superior design flexibility, and fast manufacturing speed. Here, we report a circular RNA (circRNA) vaccine that encodes the trimeric RBD of SARS-CoV-2 Spike protein. Without the need of nucleotide modification, 5’-capping or 3’-polyadenylation, circRNA could be rapidly produced via *in vitro* transcription and is highly thermostable whether stored in naked or lipid-nanoparticle (LNP)-encapsulated format. LNP-encapsulated circRNA^RBD^ elicited potent neutralizing antibodies and T cell responses, providing robust protection against Beta (B.1.351) and native viruses in mice and rhesus macaques, respectively. Notably, circRNA vaccine enabled higher and more durable antigen production than 1mΨ-modified mRNA vaccine, eliciting a higher proportion of neutralizing antibodies and stronger Th1-biased immune responses. Importantly, we found that circRNA^RBD-Omicron^ vaccine induced effective neutralizing antibodies against only Omicron but not Delta variant. By contrast, circRNA^RBD-Delta^ could elicit high level of neutralizing antibodies against both Delta and Omicron. Following two doses of either native- or Delta-specific vaccination, circRNA^RBD-Delta^, but not Omicron or Beta vaccines, could effectively boost the neutralizing antibodies against both Delta and Omicron variants. These results suggest that circRNA^RBD-Delta^ is a favorable choice for vaccination to provide a broad-spectrum protection against the current variants of concern of SARS-CoV-2.

## Main Text

Coronavirus disease 2019 (COVID-19) is a serious worldwide public health emergency caused by a novel severe acute respiratory syndrome coronavirus (SARS-CoV-2)^1,2^. So far, COVID-19 had resulted in ∼300 million confirmed cases and over 5 million confirmed deaths (World Health Organization). With the development of the epidemic, variants with immune escape ability appeared, the most serious of which is Omicron. As of Dec. 28, 2021, Omicron had taken ∼30% of COVID-19 cases (GISAID). The rapid rise of Omicron suggests that it could overtake Delta as the next dominant strain. Omicron carries over 30 mutations on the Spike protein, 15 of which are located in the receptor-binding domain (RBD)^3^, resulting in a significant decrease in the effectiveness of prior neutralizing antibodies^4–8^. This poses a severe challenge to the efficacy of current vaccines, highlighting the urgent need to develop effective vaccines against such fast-spreading variants.

SARS-CoV-2, together with the other two highly pathogenic coronaviruses, Severe Acute Respiratory Syndrome (SARS)-CoV and Middle East Respiratory Syndrome (MERS)-CoV, the other two highly pathogenic coronaviruses, belongs to the genus *Betacoronavirus* of the *Coronaviridae* family^9^. SARS-CoV-2 is a single-strand, positive-sense, enveloped virus, with an inner capsid formed by a 30-kb RNA genome wrapped by the nucleocapsid (N) proteins and a lipid envelope coated with the membrane (M), envelope (E), and trimeric Spike (S) proteins^10^. The S protein of SARS-CoV-2, composed of S1 and S2 subunits, is the major surface protein of the virion. The S protein mediates viral entry into host cells by binding to its receptor, angiotensin-converting enzyme 2 (ACE2), through the receptor-binding domain (RBD) at the C terminus of the S1 subunit. This binding subsequently induces the fusion between the SARS-CoV-2 envelope and the host cell membrane mediated by the S2 subunit, leading to the release of the viral genome into the cytoplasm^11–14^.

The S protein, S1 subunit, or the RBD antigen of SARS-CoV-2, could induce both B cell and T cell responses, generating highly potent neutralizing antibodies against SARS-CoV-2^15–17^. Vaccination is the most promising approach to end COVID-19 pandemic. Traditional vaccine platforms, such as inactivated, virus-like particle, and viral vector-based vaccines have been adopted to develop SARS-CoV-2 vaccines^18–26^. Importantly, the mRNA vaccines against SARS-CoV-2 have been developed at warp speed and urgently approved for use^27–33^, even though such a strategy had never been applied commercially before^34^. The mRNA vaccine contains a linear single-stranded RNA, consisting of a 5’ cap, the untranslated region (UTR), the antigen-coding region, and a 3’ polyA tail, and is delivered into the body via lipid-nano particle (LNP) encapsulation^34^. The clinical-scale mRNA vaccines could be manufactured rapidly upon the viral antigen sequence is released^27^. However, the current mRNA vaccine still has certain limitations, including inherent instability and suboptimal thermostability after LNP encapsulation for *in vivo* administration^35–37^, as well as potential immunogenic side effects^38,39^.

Circular RNAs (circRNAs) are covalently closed single-stranded RNA transcripts that comprise a large class of non-coding RNAs generated in eukaryotic cells by a non-canonical RNA splicing event called backsplicing in eukaryotic cells^40–42^. Some viral genomes, such as hepatitis D virus and plant viroids, are circular RNAs^39^. Thousands of circRNAs have been identified in eukaryotes, including fungi, plants, insects, fish, and mammals via high-throughput RNA sequencing and circRNA-specific bioinformatics^42^. Unlike linear mRNA, circRNA is highly stable due to its covalently closed ring structure, which protects it from exonuclease-mediated degradation^42–44^. So far, only a few endogenous circRNAs have been shown to function as protein translation templates^45–48^. Although circRNA lacks the essential elements for cap-dependent translation, it can be engineered to enable protein translation through an internal ribosome entry site (IRES) or the m6A modification incorporated to the upstream of open reading frame (ORF)^49,50^.

Here, we attempted to develop circular RNA vaccines, aiming to provide effective protection against SARS-CoV-2 and its emerging variants.

### circRNA^RBD^ produced functional SARS-CoV-2 RBD antigens

We employed the Group I ribozyme autocatalysis strategy^49^ to produce circular RNAs encoding SARS-CoV-2 RBD antigens^29^, termed circRNA^RBD^ (**Fig. 1a**). In this construct, the IRES element was placed before the RBD-coding sequence to initiate its translation. To enhance the immunogenicity of RBD antigens, the signal peptide sequence of human tissue plasminogen activator (tPA) was fused to the N-terminus of RBD to ensure the secretion of antigens^51–53^. Besides, recent research reported that RBD trimmers outperformed the monomeric RBD in binding hACE2^12,13,54^. To improve the immunogenicity of RBD antigens, the trimerization motif of bacteriophage T4 fibritin protein (foldon)^55^ was fused to its C terminus to enhance the immunogenicity of RBD antigens. This IRES-SP-RBD-Foldon sequence was then cloned into the vector to construct the *in vitro* transcription (IVT) template for producing circRNA^RBD^ (**Fig. 1a**). The precise circularization of circRNA^RBD^ was verified by reverse transcription-PCR analysis using specific primers and Sanger sequencing (**Fig. 1a**, **Extended Data Fig. 1a, b**).

**Fig. 1.**
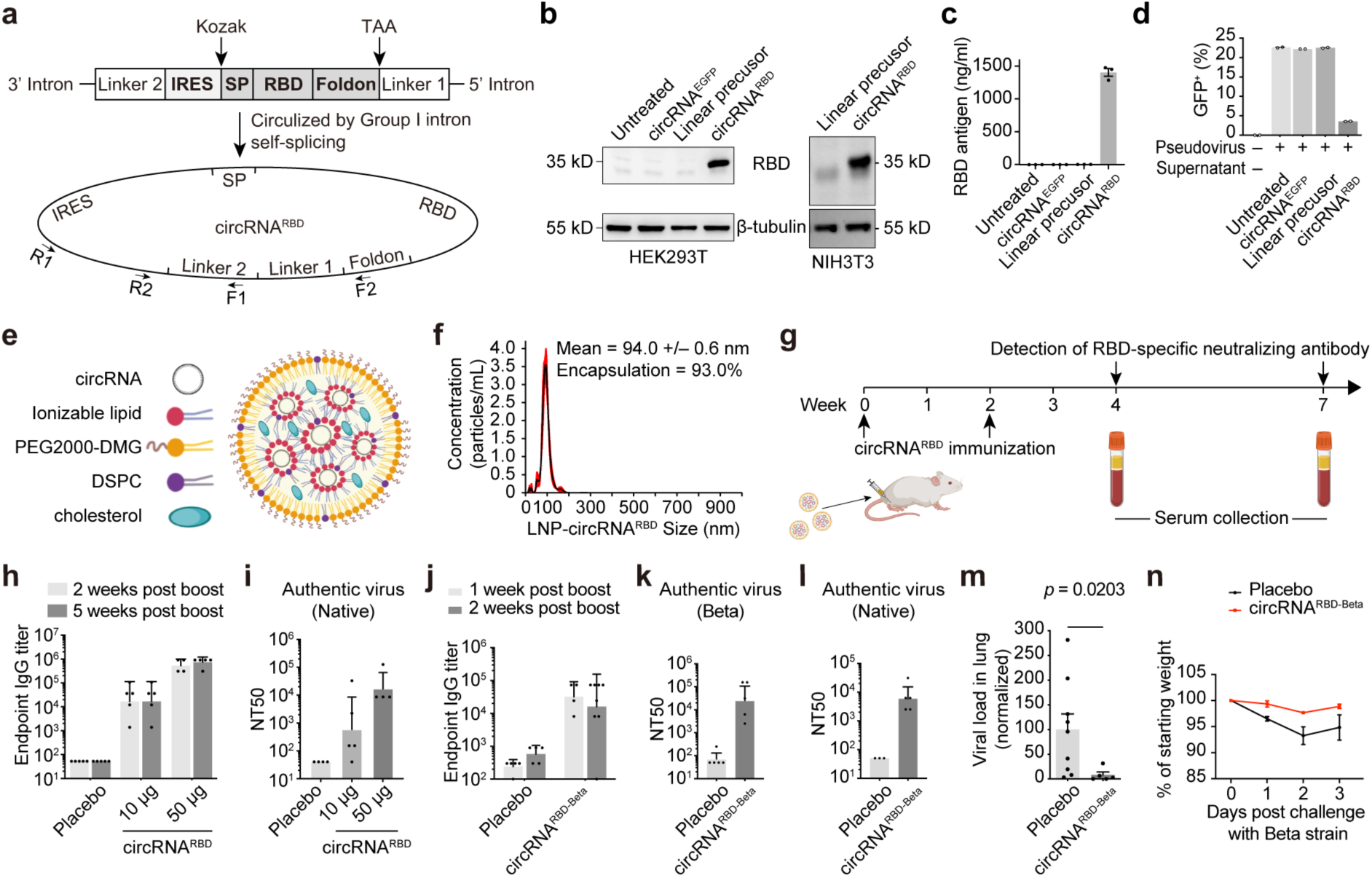
Immunogenicity and protection of circRNA vaccines against SARS-CoV-2 in mice. **a**, Schematic diagram of circRNA^RBD^ circularization by the Group I ribozyme autocatalysis. SP, signal peptide sequence of human tPA protein. Foldon, the trimerization domain from bacteriophage T4 fibritin protein. RBD, the receptor binding domain of SARS-CoV-2 Spike protein. The arrows indicate the design of primers for PCR analysis. **b**, Western Blot analysis showing the expression level of RBD antigens in the supernatant of HEK293T or NIH3T3 cells transfected with circRNA^RBD^ circularization by the Group I ribozyme autocatalysis. The circRNA^EGFP^ and linear RNA precursor were set as controls. **c**, Quantitative ELISA assay to measure the concentration of RBD antigens in the supernatant of HEK293T cells. **d**, Quantification of the competitive inhibition of SARS-CoV-2 pseudovirus infection (EGFP) by the circRNA^RBD^-translated RBD antigens in the supernatant of HEK293T cells. The circRNA^EGFP^ and linear RNA precursor were set as controls. In **c** and **d**, data were shown as the mean ± S.E.M. (n = 2 or 3). **e**, Schematic representation of LNP-circRNA complex. **f**, Representative of concentration-size graph of LNP-circRNA^RBD^ measured by dynamic light scattering method. **g**, Schematic diagram of the circRNA^RBD^ vaccination process in BALB/c mice and serum collection schedule for specific antibodies analysis. **h**, Measuring the SARS-CoV-2 specific IgG antibody endpoint GMTs elicited by circRNA^RBD^ vaccine with ELISA. **i**, Neutralization assay of SARS-CoV-2 authentic virus with the sera of mice immunized with circRNA^RBD^ vaccines. The serum samples were collected at 5 weeks post boost. **j**, Measuring the SARS-CoV-2 (Beta) specific IgG antibody endpoint GMTs elicited by circRNA^RBD-Beta^ vaccine with ELISA. **k**, Neutralization assay of SARS-CoV-2 (Beta) authentic virus with the serum of mice immunized with circRNA^RBD-Beta^ vaccines. **l**, Neutralization assay of SARS-CoV-2 (D614G) authentic virus with the serum of mice immunized with circRNA^RBD-Beta^ vaccines. In **h** and **l**, data were shown as the geometric mean ± geometric S.D., and each symbol represented an individual mouse. **m**, Viral loads in the lung tissues of mice challenged with authentic SARS-CoV-2 Beta. In **m**, data were shown as the mean ± S.E.M. (n >= 5), and the statistical test was performed by unpaired two-sided Student’s *t*-test. **n**, The weight change of immunized or placebo mice after SARS-CoV-2 Beta challenge. In **n**, data were shown as the mean ± S.E.M. (n >= 5).

Owing to its covalently closed circular structure, the circRNA^RBD^ migrated faster in electrophoresis (**Extended Data Fig. 2a**) and appeared more resistant to exonuclease RNase R than the linear precursor RNA (**Extended Data Fig. 2b**). High-performance liquid chromatography (HPLC) showed that the RNase R treatment purged a significant amount of the linear precursor RNAs, which was an important step in purifying the circRNA^RBD^ (**Extended Data Fig. 2c**).

To test the secretory expression of RBD antigens produced by circRNA^RBD^, the purified circRNA^RBD^ was transfected into human HEK293T cells or murine NIH3T3 cells. Abundant RBD antigens in the supernatant of both human and murine cells was detected by Western blot, indicating the high compatibility of circRNAs (**Fig. 1b**). The concentration of RBD antigens produced by circRNA^RBD^ reached ∼1,400 ng/mL, 600-fold higher than its linear precursor RNA (**Fig. 1c**).

Besides the Group I ribozyme autocatalysis strategy, we developed an alternative method to generate circRNA^RBD^ using T4 RNA ligase (**Extended Data Fig. 3a**). Similarly, abundant RBD antigens have been detected in the supernatant at a concentration of ∼1,000 ng/mL concentration, which is ∼200-fold higher than that produced by its linear precursor RNA (**Extended Data Fig. 3b, c**).

To verify whether the secreted SARS-CoV-2 RBD antigens produced by circRNA^RBD^ were functional, the supernatants of circRNA^RBD^-transfected cells were used in a competition assay using hACE2-overexpressing HEK293 cells (HEK293T-ACE2) and SARS-CoV-2 pseudovirus harboring an EGFP reporter^56^. The secreted SARS-CoV-2 RBD antigens could effectively block SARS-CoV-2 pseudovirus infection (**Fig. 1d**). Altogether, circRNA^RBD^ showed robust protein expression, suggesting its potential as a novel vaccine platform.

### SARS-CoV-2 circRNA^RBD^ vaccine induced sustained humoral immune responses with high-level of neutralizing antibodies

To explore whether circRNA could be leveraged to create a new type of vaccine, we attempted to assess the immunogenicity of circRNA^RBD^ encapsulated with lipid nanoparticles in BALB/c mice (**Fig. 1e**). The circRNA^RBD^ encapsulation efficiency was greater than 93%, with an average diameter of 100 nm (**Fig. 1f**). Mice were immunized with LNP-circRNA^RBD^ through intramuscular (i.m.) injection twice at a two-week interval, at doses of 10 µg or 50 µg per mouse, with empty LNP serving as the placebo (**Fig. 1g**). The amount of RBD-specific binding IgG and neutralizing antibodies was measured at two- or five-weeks post the boost dose. The circRNA^RBD^ elicited high level of RBD-specific IgG endpoint geometric mean titers (GMTs), reaching ∼1.9×10^4^ for 10 µg dose and ∼5.7×10^5^ for 50 µg dose, indicating that circRNA^RBD^ could induce SARS-CoV-2 RBD-specific antibodies (**Fig. 1h**).

Sera from circRNA^RBD^-vaccinated mice could effectively neutralize both SARS-CoV-2 pseudovirus with 50% neutralization titer (NT50) of ∼4.5×10^3^ (**Extended Data Fig. 4**) and authentic SARS-CoV-2 virus with NT50 of ∼7.0×10^4^ (**Fig. 1i**). The high level of RBD-specific IgG, potent RBD antigen neutralization, and sustained SARS-CoV-2 neutralizing capacity confirmed that circRNA^RBD^ vaccines induced humoral immune responses in mice.

### SARS-CoV-2 circRNA^RBD-Beta^ vaccine-elicited antibodies showed preferential neutralizing activity against Beta variant

Next, we evaluated the efficacy of a circRNA^RBD-Beta^, a circRNA vaccine encoding RBD/K417N-E484K-N501Y antigens derived from SARS-CoV-2 Beta variant. BALB/c mice were immunized with an i.m. injection of the circRNA^RBD-Beta^ vaccine, followed by a boost at a two-week interval. The immunized mice’s sera were collected at 1 and 2 weeks post the boost. ELISA showed that the RBD-Beta-specific IgG endpoint GMT was ∼1.6×10^4^ at two weeks post boost (**Fig. 1j**). SARS-CoV-2 pseudovirus neutralization assay revealed that circRNA^RBD^-elicited antibodies could effectively neutralize all three pseudoviruses, with the highest neutralizing activity against the native (D614G) strain (**Extended Data Fig. 5a**). The circRNA^RBD-Beta^-elicited antibodies could also neutralize all three pseudoviruses, with the highest activity against its corresponding Beta variant (**Extended Data Fig. 5b**).

In line with pseudovirus neutralization assay, the sera from immunized mice could neutralize authentic SARS-CoV-2 Beta and native (D164G) strain with NT50 of 2.6×10^4^ (**Fig. 1k**) and 6.0×10^3^ (**Fig. 1l**), respectively. Collectively, circRNA vaccine-elicited antibodies showed the highest neutralizing activity against their corresponding variant strains. It’s worth noting that both vaccines could neutralize all three strains, albeit with varying efficacies. Nevertheless, it’s likely that the updated vaccines based on the mutations of the emerging variants might provide better and more effective protection against SARS-CoV-2 circulating variants.

### circRNA^RBD-Beta^ vaccine protected mice against the infection of Beta variant

To further evaluate the protective efficacy of circRNA^RBD-Beta^ vaccine *in vivo*, we employed the Beta variant for authentic virus challenge experiments. Consistent with a recent report^57,58^, the Beta variant could infect wild-type BALB/c mice and replicate in their lungs (**Extended Data Fig. 5c**), likely due to the mutations in Spike protein such as K417N, E484Q, and N501Y^57,59^. Seven weeks post the boost dose, the RBD-Beta-specific IgG endpoint GMT was still around 1.2×10^4^ (**Extended Data Fig. 5d**), with significant neutralizing activity against RBD-Beta antigens (**Extended Data Fig. 5e**). Each immunized mouse was then intranasally infected with 5×10^4^ PFU of Beta virus (7 weeks post the boost dose). The lung tissues were collected three days after the challenge for the detection of viral RNAs. Viral loads in the lungs of vaccinated mice were significantly lower compared with the placebo group (**Fig. 1m**). Consistently, only the mice in the placebo group underwent weight loss (**Fig. 1n**). These results indicated that the circRNA^RBD-Beta^ vaccine could effectively protect the mice against SARS-CoV-2 Beta.

Considering that the 50 µg of circRNA^RBD^ elicited a higher level of neutralizing antibodies than 10 µg, we postulated that the LNP delivery might have great impact on the efficacy of circular RNA vaccine. After multiple tests, we were able to significantly lower the vaccine dose using one of the commercial formulas (Precision Nanosystems). 10 µg of circRNA could induce neutralizing antibodies at a level comparable to 50 µg (**Extended Data Fig. 6**). We thus switched our choice of LNP for the rest of our experiments.

### circRNA^RBD-Delta^ vaccine induced potent neutralizing antibodies against SARS-CoV-2 Delta

The Delta, like the Beta variant, partially escapes the antibodies produced in the survivors or vaccinees^60–62^. To develop such a variant-specific vaccine, we adopted both Group I ribozyme autocatalysis and T4 RNA ligation strategies to produce circRNA^RBD-Delta^. Mice were immunized i.m. with 0.5 µg, 2.5 µg, 5 µg or 10 µg of circRNA^RBD-Delta^ vaccines twice at a two-week interval. Two weeks post the boost dose, the sera from immunized mice were collected to detect RBD-specific antibodies. Vaccines of circRNA^RBD-Delta^ made by either circularization methods could induce high endpoint GMTs of IgG binding antibodies (**Fig. 2a, b**). The sera from circRNA^RBD-Delta^-vaccinated mice effectively neutralized the Delta pseudovirus in a dose-dependent manner, with the NT50 of ∼1.4×10^5^ for the 10 µg dose (**Fig. 2c**).

**Fig. 2.**
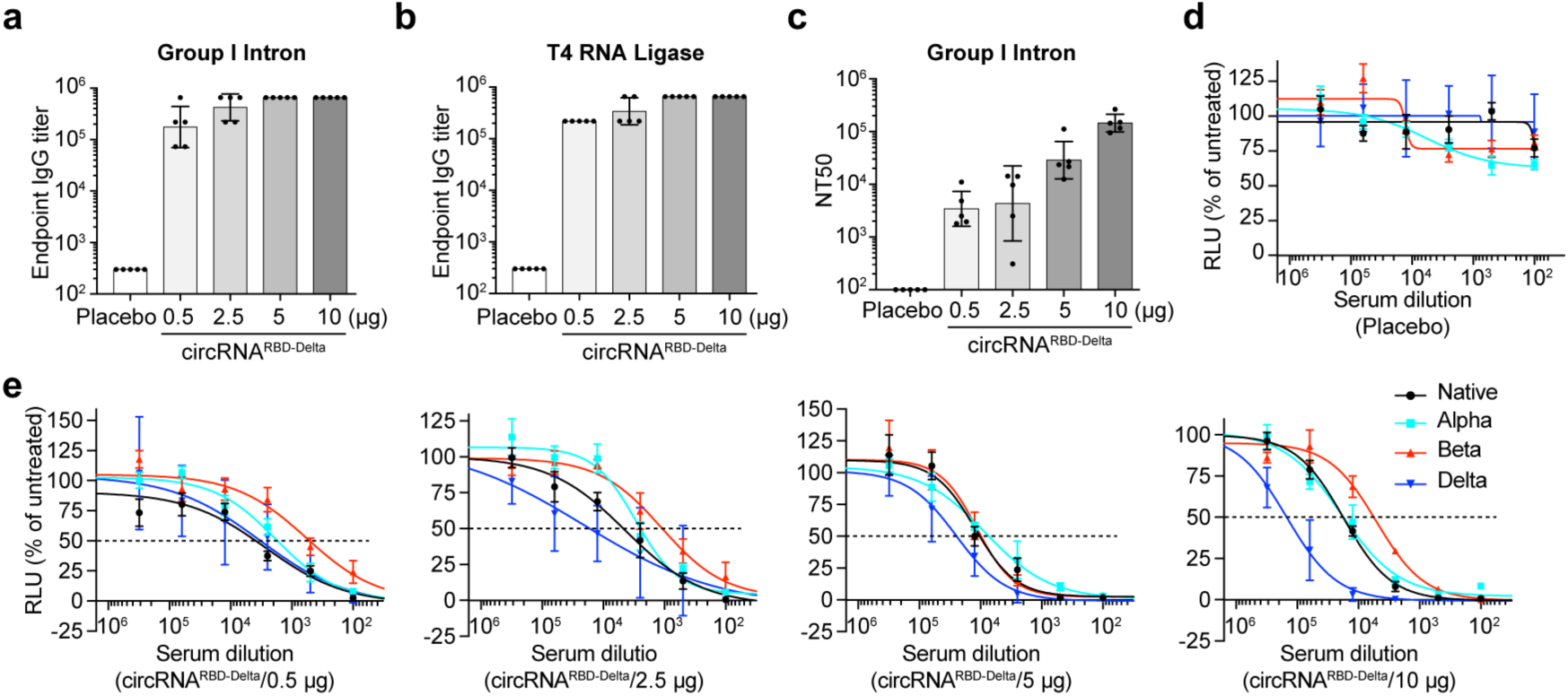
Humoral immune responses elicited by circRNA^RBD-Delta^ vaccines in mice. **a**, Measuring the SARS-CoV-2 Delta specific IgG antibody endpoint GMTs of circRNA^RBD-Delta^ vaccines generated by Group I ribozyme autocatalysis with ELISA. The data were shown as the mean ± S.E.M. (n = 5). **b**, Measuring the SARS-CoV-2 Delta specific IgG antibody endpoint GMTs of circRNA^RBD-Delta^ vaccines generated by T4 RNA ligase with ELISA. The data were shown as the mean ± S.E.M. (n = 5). **c**, Neutralization assay of VSV-based SARS-CoV-2 (Delta) pseudovirus with the sera of mice immunized with circRNA^RBD-Delta^ vaccines. The data were shown as the mean ± S.E.M. (n = 5). In **a**-**c**, data were shown as the geometric mean ± geometric S.D., and each symbol represented an individual mouse. **d** and **e**, Neutralization assay of SARS-CoV-2 native, Alpha, Beta and Delta pseudovirus with the sera of mice immunized with placebo control (**d**), or 0.5 µg, 2.5 µg, 5 µg, or 10 µg of circRNA^RBD-Delta^ vaccines (**e**). In **d** and **e**, data were shown as the mean ± S.E.M. (n >= 5). In **a**-**e,** the serum samples were collected at 2 weeks post boost.

Importantly, circRNA^RBD-Delta^ vaccines could provide protection against other variants, including the native strain, Alpha and Beta variants, albeit with varying degrees. The sera from circRNA^RBD-Delta^-immunized mice exhibited the highest neutralizing activity against Delta and the lowest against Beta variant (**Fig. 2d, e**).

### circRNA vaccine enabled higher and more durable antigen expression than mRNA vaccine

The circRNA is reportedly more stable than mRNA owing to its covalent closed circular structure^63^. To test whether the stability of the circRNA vaccine could confer higher and more durable antigen-encoding efficiency than the mRNA vaccine, we generated 1mΨ modified mRNA, termed 1mΨ-mRNA, and unmodified mRNA, both of which contain the same RBD-encoding sequence as the circRNA for a fair comparison. The purified circRNA, 1mΨ-mRNA, and unmodified mRNA were transfected into the HEK239T cells. Cell supernatants were collected to measure the abundance of RBD antigen at different time points from 12 to 96 hrs post transfection. The circRNA produced much higher levels of RBD antigens at all time points and maintained for longer period than both 1mΨ-mRNA and unmodified mRNA (**Fig. 3a**). The RT-qPCR showed that circRNAs were more stable than mRNAs, modified or unmodified (**Fig. 3b**). Importantly, the LNP-encapsulation further enhanced the advantage of circRNA in protein production and durability from both 1mΨ-mRNA and unmodified-mRNA (**Fig. 3c**). Interestingly, the LNP-encapsulation appeared to improve the antigen-encoding efficiency of unmodified mRNA to a level comparable to 1mΨ-mRNA (**Fig. 3c**).

**Fig. 3.**
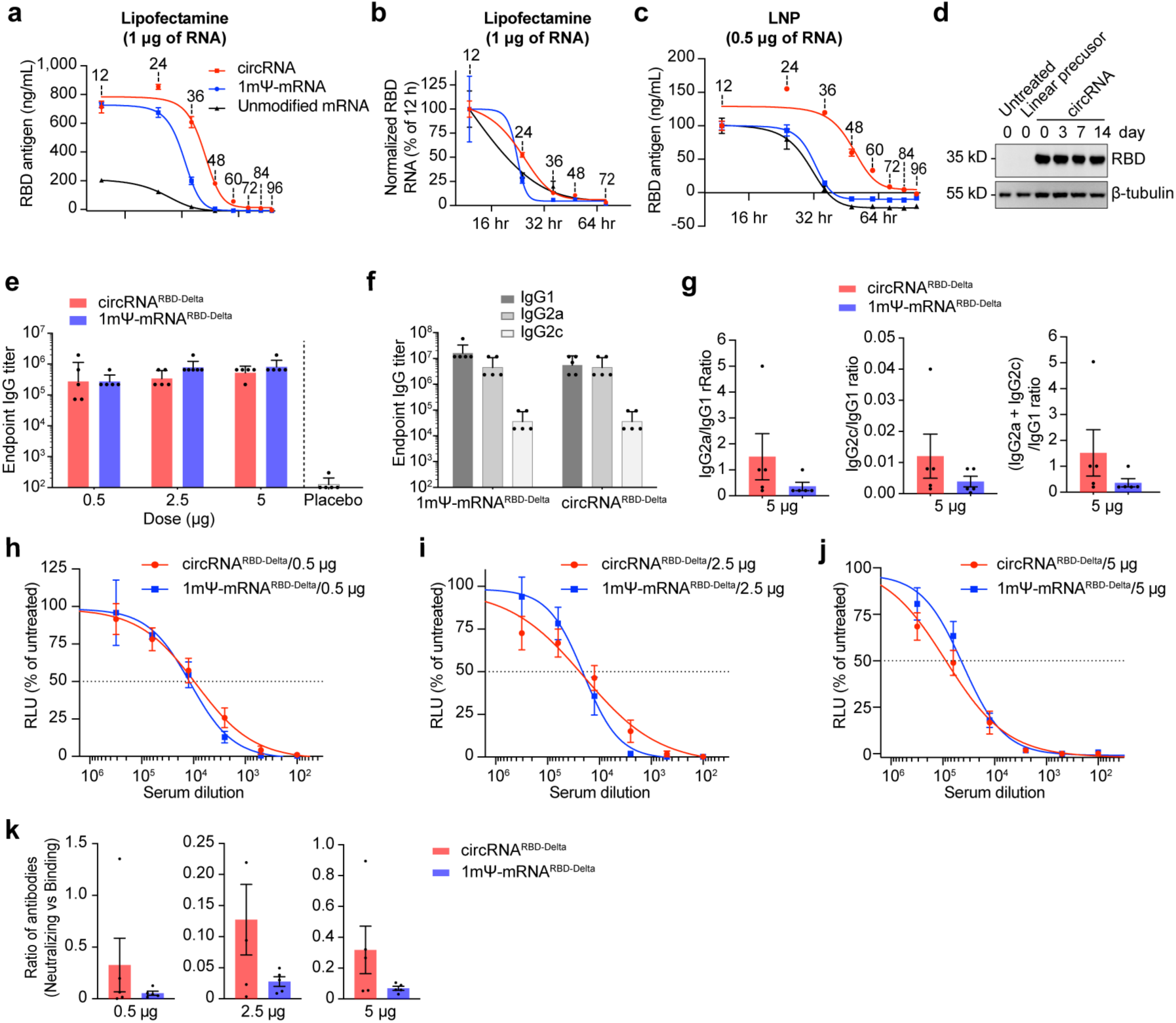
circRNA vaccine elicited higher proportion of neutralizing antibodies and stronger Th1-biased T cell immune responses than mRNA vaccine. **a**, Comparing antigen expression level of circRNA^RBD-Delta^, 1mΨ-mRNA^RBD-Delta^ and non-modified mRNA^RBD-Delta^ through lipofectamine transfection in HEK293T cells by ELISA at different time points. **b**, The dynamic change of RNA level in (**a**). **c**, Comparing antigen expression level of LNP-circRNA, LNP-1mΨ-mRNA or LNP-non-modified-mRNA delivered into HEK293T cells with ELISA at indicated time points. In **a**-**c**, data were shown as the mean ± S.E.M. (n = 3). The data were shown as the mean ± S.E.M. (n = 3). **d**, Western Blot analysis showing the expression level of RBD antigens in the supernatant of HEK293T cells transfected with circRNA^RBD^. The circRNA^RBD^ were stored at room temperature and transfected in days 0, 3, 7 and 14. The untreated and linear precursor were used as controls. **e**, Measuring the RBD-Delta-specific IgG endpoint GMTs elicited by circRNA^RBD^ vaccine or mRNA vaccine. The data were shown as the mean ± S.E.M. (n = 5). **f**, Measuring RBD-Delta-specific IgG1/IgG2a/IgG2c endpoint GMTs elicited by 5 µg of circRNA^RBD^ vaccine or mRNA vaccine in mice. In **e** and **f**, data were shown as the geometric mean ± geometric S.D., and each symbol represented an individual mouse. **g**, Measuring the specific IgG2a/IgG1, IgG2c/IgG1 and (IgG2a + IgG2c)/IgG1 ratio of serum from mice immunized with 5 µg of circRNA^RBD-Delta^ or mRNA^RBD-Delta^. **h**-**j**, Sigmoidal curve diagram of neutralization rate of VSV-based SARS-CoV-2 (Delta) pseudovirus with the sera from mice immunized with different dose of circRNA^RBD-Delta^ or mRNA^RBD-Delta^ vaccines. The serum samples were collected at 2 weeks post boost. **k**, Measuring the ratio of (Neutralizing antibodies)/(Binding antibodies) elicited by circRNA^RBD-Delta^ or mRNA^RBD-Delta^ vaccine. The ratio of (NT50)/(Endpoint GMT) of each mouse was calculated. In **g**-**k**, data were shown as the mean ± S.E.M. (n >= 4) and each symbol represented an individual mouse.

A thermostable RNA vaccine is highly desirable for efficient vaccine distribution. We found that even after two weeks of storage at room temperature (∼25°C), the circRNA could express RBD antigens without detectable loss (**Fig. 3d**), highlighting its remarkable thermal stability. To further evaluate the thermostability of vaccines, the LNP-encapsulated circRNA, 1mΨ-mRNA and unmodified-mRNA were stored at 4°C, room temperature (∼25°C), or 37°C for up to 28 days prior to the transfection. At all temperatures tested, circRNA expressed higher levels of antigens than those of the other two mRNA groups (**Extended Data Fig. 7a-c**). At 4°C, little reduction of RBD antigens produced by LNP-encapsulated circRNA could be detected from 1-28 days (**Extended Data Fig. 7a**). The stability of LNP-encapsulated circRNA, 1mΨ-mRNA and unmodified-mRNA was evidently reduced with the increase of storage temperature, especially at 37°C (**Extended Data Fig. 7b-c**). Together, although extended shelf life and high temperature deteriorate RNA-based vaccines, circRNA vaccine exhibited higher stability than mRNA vaccines (**Extended Data Fig. 7a-c**).

### circRNA vaccine elicited higher proportion of neutralizing antibodies and stronger Th1-biased T cell immune responses than the mRNA vaccine

Currently, two kinds of mRNA vaccines are widely inoculated, mRNA-1273 (Moderna) and BNT162b2 (Pfizer/BioNTech), both of which contain modified 1mΨ modification. Given that circRNA vaccine possesses higher stability and antigen-encoding efficiency, we wonder whether it exhibited superior immunogenicity to mRNA vaccine. We first compared the balance of Th1 and Th2 immune responses between circRNA^RBD-Delta^ and mRNA^RBD-Delta^ vaccines because Th2-biased immune responses might induce vaccine-associated enhanced respiratory disease (VAERD)^27,32,64^. ELISA assay showed that the total IgG elicited by circRNA^RBD-Delta^ was comparable to that by mRNA^RBD-Delta^ (**Fig. 3e**), however, the ratio of IgG2a/IgG1, IgG2c/IgG1 or (IgG2a + IgG2c)/IgG1 from circRNA^RBD-Delta^ was higher than from mRNA^RBD-Delta^ vaccine (**Fig. 3f, g**, **Extended Data Fig. 8a, b**), indicating that circular RNA vaccines tended to induce stronger Th1-biased immune responses.

Antibody-mediated enhancement (ADE) of infection by virus-specific antibodies is another potential concern for vaccines, which has been reported for infections by some viruses including Zika, Dengue virus, and coronaviruses^65–69^. Previous research has reported that the virus-binding antibodies without neutralizing activity elicited by infection or vaccination possibly caused the ADE effects, especially for those viruses with different serotypes^70,71^. Therefore, we compared the ratio of neutralizing to binding antibodies between circRNA and mRNA vaccines. Although circRNA^RBD-Delta^ exhibited equal neutralizing capability compared to mRNA^RBD-Delta^ (**Fig. 3h-j**), the former induced a higher proportion of neutralizing antibodies at all doses tested (0.5 µg, 2.5 µg, and 5 µg) in mice (**Fig. 3k**). Owing to this unique feature, the circRNA vaccine might have certain advantage to circumvent potential ADE effects and tolerate viral mutations.

### circRNA^RBD-Delta^ vaccine elicited strong T cell immune responses

B cells, CD4^+^ T cells, and CD8^+^ T cells are three pillars of adaptive immunity. They mediated effector functions against SARS-CoV-2 in both non-hospitalized and hospitalized cases of COVID-19^72^.

To compare CD4^+^ and CD8^+^ T cell immune responses, the splenocytes of immunized mice were collected and stimulated with SARS-CoV-2 RBD-Delta pooled peptides (**Supplementary Table 1**), and cytokine-producing T cells were quantified by intracellular cytokine staining among effector memory T cells (Tem, CD44^+^CD62L^-^) (**Extended Data Fig. 9**). Stimulated with RBD-Delta peptide pools, CD8^+^ T cells producing interferon-γ (IFN-γ), tumor necrosis factor (TNF-α), and interleukin-2 (IL-2) were detected in mice immunized with circRNA^RBD-Delta^ vaccine or 1mΨU-mRNA^RBD-Delta^ vaccine (**Extended Data Fig. 10a-c**), indicating the RBD-specific CD8^+^ T cell responses elicited by both vaccines. The CD4^+^ T cells of immunized mice induced strong IFN-γ, TNF-α, and IL-2 responses, but minimal IL-4 responses (**Extended Data Fig. 10d-g**), indicating that both circRNA^RBD-Delta^ and 1mΨ-mRNA^RBD-Delta^ induced the Th1-biased T cell immune responses (**Extended Data Fig. 10d-g**).

### circRNA^RBD-Delta^ vaccine elicited high level of broad-spectrum neutralizing antibodies against both Delta and Omicron variants

The emerging SARS-CoV-2 Omicron variant contains more than 30 mutations in the Spike region and spreads quickly^3^. It has been reported that the Omicron variant escaped most of the previously reported neutralizing antibodies and the sera of vaccinees who received two-dose original SARS-CoV-2 vaccines^4–8^. To cope with this emergency, we tested the neutralizing capability elicited by all three circRNA vaccines against Omicron variant. The neutralizing activity against Omicron from the sera of mice immunized with either one of the three vaccines dropped 74-fold (native), 15-fold (Beta) and 44-fold (Delta) in comparison with against their corresponding variants (**Fig. 4a**). Among all three, circRNA^RBD-Delta^ vaccine maintained sufficient neutralizing activity against Omicron (**Fig. 4a**), with the NT50 of ∼4.7×10^3^, while the NT50 of the circRNA^RBD-Beta^ against Omicron dropped to below 5×10^2^ (**Fig. 4a**). Compared to mRNA^RBD-Delta^ vaccine, circRNA^RBD-Delta^ vaccine elicited comparable neutralizing activity against both Delta and Omicron variants for mouse sera collected at 2 weeks post boost (short-term) and 7 weeks post boost (long-term) (**Fig. 4a-c**). Similar to the above observations (**Fig. 3h-k**), circRNA^RBD-Delta^ vaccine also elicited higher proportion of neutralizing antibodies against Omicron variant than 1mΨ-mRNA^RBD-Delta^ vaccine at both 2 weeks post boost (short-term) and 7 weeks post boost (long-term) (**Extended Data Fig. 11a-d**), indicating the potential superiority of circRNA vaccine against the circulating variants of SARS-CoV-2, likely owing to its high proportion of neutralizing antibody (**Fig. 3h-k** and **Extended Data Fig. 11a-d**).

**Fig. 4.**
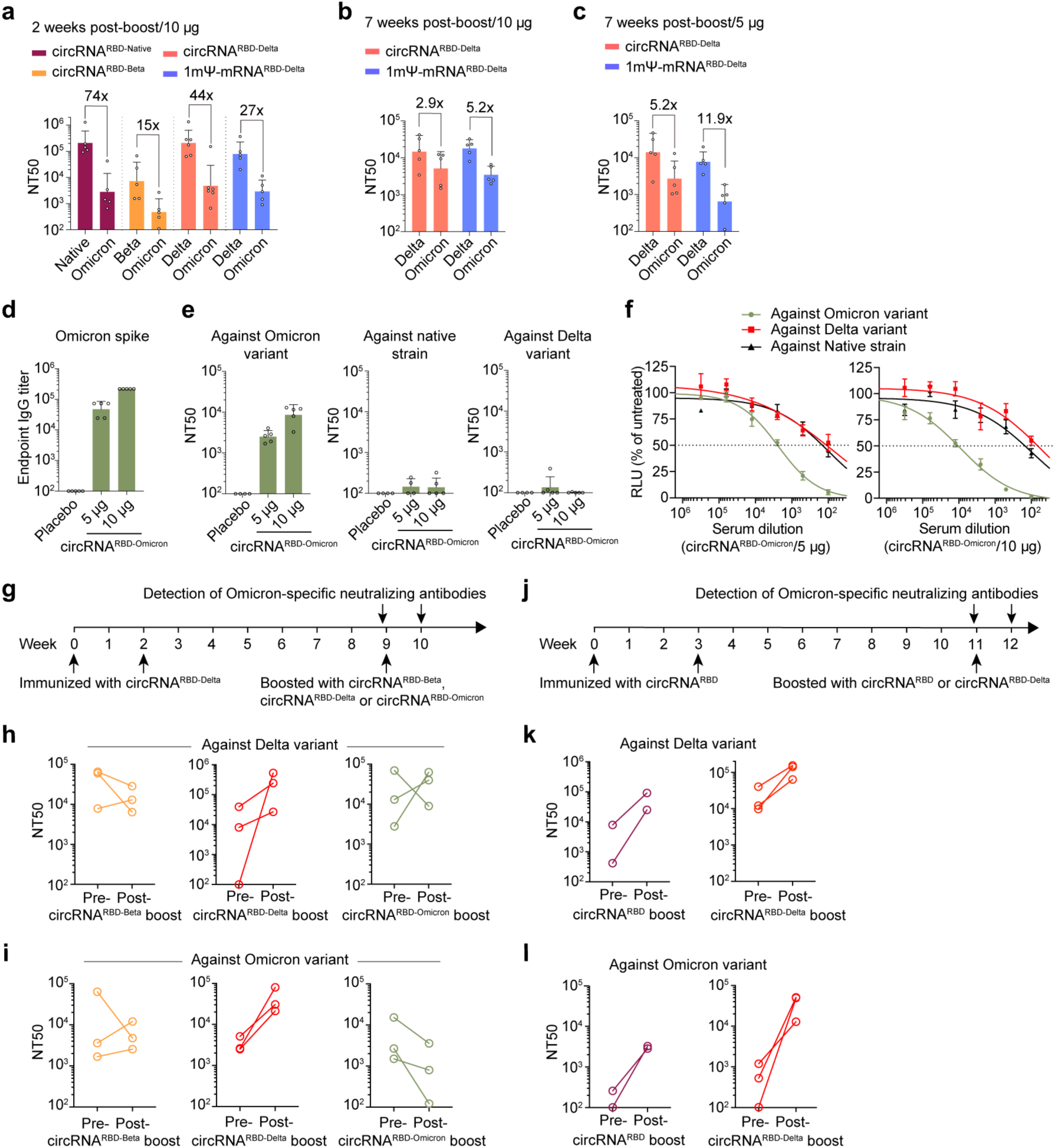
circRNA^RBD-Delta^ vaccine elicited high-level of neutralizing antibodies against both Delta and Omicron variants. **a**, Neutralization assay of VSV-based SARS-CoV-2 pseudovirus with the sera of mice immunized with 10 µg of circRNA^RBD^, circRNA^RBD-Beta^, circRNA^RBD-Delta^ or mRNA^RBD-Delta^ vaccines. The serum samples were collected at two weeks post boost. **b** and **c**, Neutralization assay of VSV-based SARS-CoV-2 pseudovirus with the sera of mice immunized with 10 µg (**b**) or 5 µg (**c**) of circRNA^RBD-Delta^ or mRNA^RBD-Delta^ vaccines. The serum samples were collected at 7 weeks post boost. **d**, Measuring the Omicron-Spike-specific IgG endpoint GMTs of circRNA^RBD-Omicron^-immunized mouse sera with ELISA. **e**, Measuring the NT50 of LNP-circRNA^RBD-Omicron^-immunized mouse sera using VSV-based pseudovirus of Omicron, Delta and native strain. The serum samples were collected at 1 week post boost dose. In **a**-**e**, data were shown as the geometric mean ± geometric S.D., and each symbol represented an individual mouse. **f**, Sigmoidal curve diagram of the neutralization assay using VSV-based pseudovirus of Omicron, Delta and native strain with the sera of mice immunized with 5 µg or 10 µg of circRNA^RBD-Omicron^ vaccines. In **f**, the serum samples were collected at 1 week post boost, and data were shown as the mean ± S.E.M. (n = 4 or 5). g, Schematic diagram of circRNA vaccination process in BALB/c mice and serum collection schedule for detecting specific neutralizing antibodies. **h** and **i**, Measuring the NT50 value of 5 µg of circRNA^RBD-Beta^, circRNA^RBD-Delta^ or circRNA^RBD-Omicron^-boosted mouse sera after receiving two-dose circRNA^RBD-Delta^ vaccine (5 µg) using VSV-based pseudovirus of Delta (**h**) or Omicron (**i**). In **h** and **i**, each symbol represented an individual mouse. **j**, Schematic diagram of the LNP-circRNA vaccination process in BALB/c mice and serum collection schedule for specific antibodies analysis. **k** and **l**, Measuring the NT50 value of mouse sera boosted with 20 µg of circRNA^RBD^ or circRNA^RBD-Delta^ vaccine after receiving two-dose circRNA^RBD^ vaccine (20 µg) using VSV-based pseudovirus of Delta (**k**) or Omicron (**l**). In **k** and **l**, each symbol represented an individual mouse.

### circRNA^RBD-Omicron^ vaccine elicited neutralizing antibodies against Omicron

We have developed Omicron-specific circRNA vaccine, which expressed the trimeric RBD antigens of Omicron variant. Mice were immunized i.m. with 5 µg or 10 µg of circRNA^RBD-Omicron^ vaccines twice at a 2-week interval. One week post the boost dose, the serum samples from immunized mice were collected for the detection of Omicron-specific antibodies. The circRNA^RBD-Omicron^ vaccine could induce Omicron Spike-specific antibodies with the endpoint GMTs of ∼4.7×10^4^ for 5 µg dose and ∼2.2×10^5^ for 10 µg dose (**Fig. 4d**), yielding evident neutralizing activities against Omicron with the NT50 of ∼2.5×10^3^ for the 5 µg dose and ∼8.6×10^3^ for the 10 µg dose (**Fig. 4e**). However, neutralizing activity could hardly be detected against native strain or Delta variant (**Fig. 4e, f**).

### The third booster with circRNA^RBD-Delta^ vaccine remarkably elevated the neutralizing antibodies against current VOC of SARS-CoV-2

We next investigated the feasibility of circRNA vaccine as a booster for the protection against Omicron variant. Mice immunized with two doses of circRNA^RBD-Delta^ vaccines received the 3^rd^ booster with circRNA^RBD-Beta^, circRNA^RBD-Delta^ or circRNA^RBD-Omicron^ vaccine at 7 weeks post the 2^nd^ dose, followed by the assessment of neutralizing activity against the Omicron variant at 1 week post the 3^rd^ boost (**Fig. 4g**). Only circRNA^RBD-Delta^ effectively boosted the neutralizing antibodies against both Delta (**Fig. 4h**) and Omicron (**Fig. 4i**). On the contrary, the 3^rd^ boost with the circRNA^RBD-Beta^ or circRNA^RBD-Omicron^ vaccine failed to elevate the neutralizing capability against Omicron (**Fig. 4h, i**).

We then went on to test the 3^rd^ booster with circRNA^RBD^ or circRNA^RBD-Delta^ vaccine on mice previously immunized with 2 doses of circRNA^RBD^ vaccines (**Fig. 4j**). Both vaccines could effectively boost the neutralizing antibodies against both Delta (**Fig. 4k**) and Omicron (**Fig. 4l**). circRNA^RBD-Delta^ appeared to be a much better booster than circRNA^RBD^ against both Delta and Omicron variants, which elevated the geometric mean NT50 from ∼4×10^2^ to ∼3.2×10^4^ against Omicron variant (**Fig. 4k, l**).

Taken together, these results suggest that circRNA^RBD-Delta^ might be a favorable choice for vaccination to provide a broad-spectrum protection against the current VOC of SARS-CoV-2. However, the 3^rd^ booster with Omicron-specific vaccines might not be an appropriate strategy against the current Delta and Omicron emergency in the real world.

### circRNA vaccine elicited potent neutralizing antibodies and Th1-biased immune responses in rhesus macaques

To further assess the immunogenicity of circRNA vaccine in non-human primates (NHPs), groups of 2∼4-year-old rhesus macaques were immunized i.m. with 20 µg, 100 µg or 500 µg of circRNA^RBD^ vaccines, 100 µg of circRNA^Ctrl^, or PBS control on Days 0 and 21 (**Fig. 5a**). The total RBD-specific IgG binding and neutralizing antibodies were measured using the plasma of rhesus macaques at two weeks post the boost dose (**Fig. 5a**). The IgG endpoint GMTs reached ∼2.1×10^4^ (20 µg), ∼1.6×10^4^ (100 µg dose) and ∼7×10^3^ (500 µg dose) for circRNA^RBD^ vaccines, while circRNA^Ctrl^ or PBS-immunized rhesus macaques failed to induce RBD-specific antibodies (**Fig. 5b**). The SARS-CoV-2 pseudovirus neutralization assay showed the NT50 of ∼180 for 20 µg dose, ∼520 for 100 µg dose, and ∼390 for 500 µg dose (**Fig. 5c**). The authentic SARS-CoV-2 neutralization assay showed the NT50 of ∼80 for 20 µg dose, ∼120 for 100 µg dose, and ∼50 for 500 µg dose (**Fig. 5d, e**). These results indicated that the 100 µg dose could elicit maximal level of neutralizing antibodies (**Fig. 5c, d**).

**Fig. 5.**
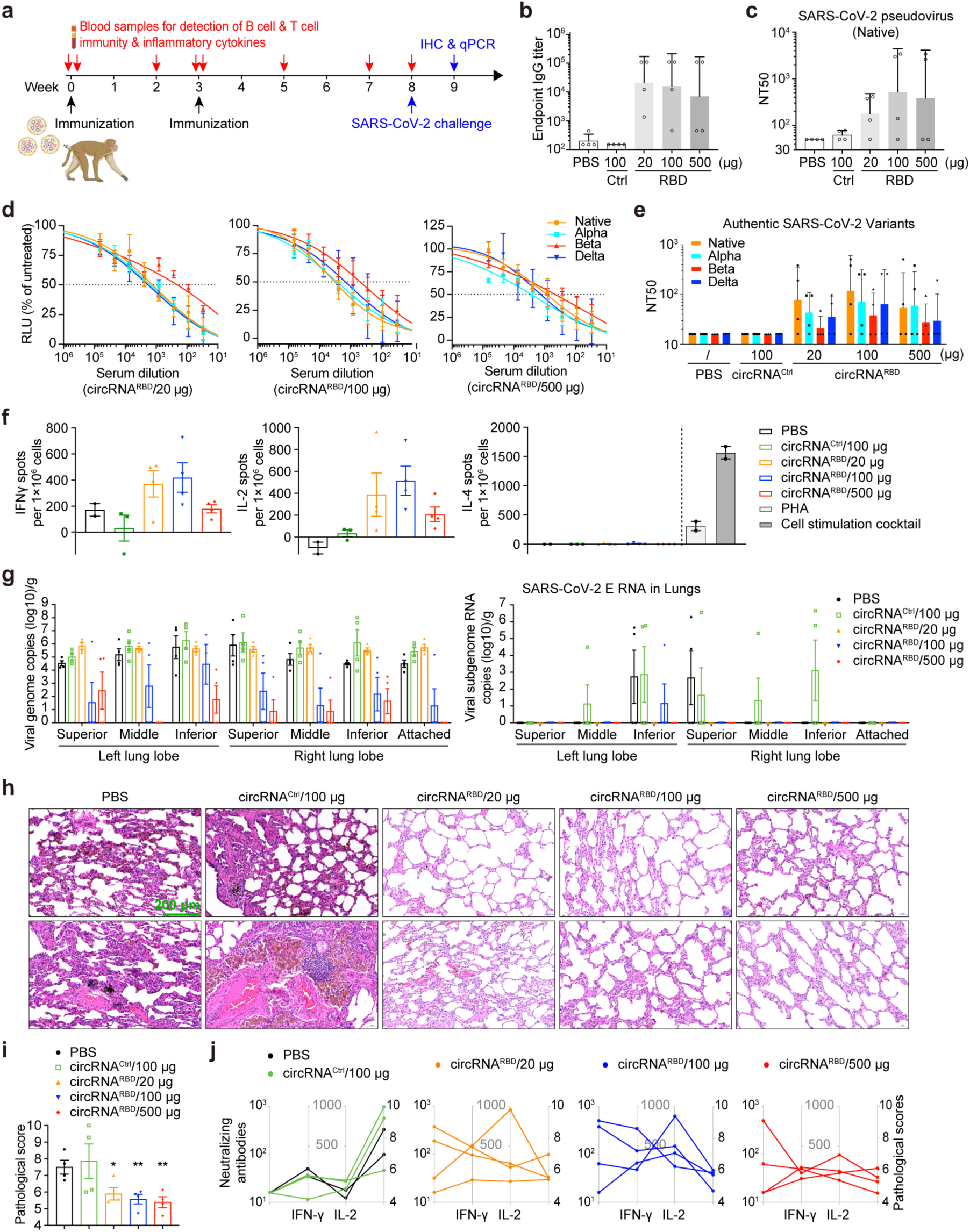
circRNA vaccine-elicited immunogenicity and protection against SARS-CoV-2 infection in rhesus macaques. **a**, Schematic diagram of the circRNA^RBD^ vaccination process in rhesus macaques and plasma collection schedule for specific antibodies analysis. **b**, Measuring the SARS-CoV-2 RBD-specific IgG endpoint GMTs of the plasma from the rhesus macaques immunized with circRNA^RBD^ vaccine, or circRNA^Ctrl^ (circRNA without the RBD-encoding sequence) or PBS controls using ELISA assay. The data were shown as the mean ± S.E.M. (n = 4). **c**, Measuring the NT50 of the plasma from the rhesus macaques immunized with circRNA^RBD^ vaccine with ELISA using VSV-based SARS-CoV-2 pseudovirus. In **b** and **c**, data were shown as the geometric mean ± geometric S.D., and each symbol represented an individual mouse. **d**, Sigmoidal curve diagram of neutralization rate of VSV-based SARS-CoV-2 native, Alpha, Beta and Delta pseudovirus using the plasma from the rhesus macaques immunized with circRNA^RBD^ vaccine. The data were shown as the mean ± S.E.M. (n = 4), and each symbol represented an individual rhesus macaque. **e**. Neutralization assay of authentic SARS-CoV-2 native, Alpha, Beta and Delta virus using the plasma from the rhesus macaques immunized with circRNA^RBD^ vaccine. In **e**, data were shown as the geometric mean ± geometric S.D., and each symbol represented an individual mouse. The data were shown as the mean ± S.E.M. (n = 4). **f**, Measuring the SARS-CoV-2 RBD-specific IFN-γ, IL-2 and IL-4 responses of PBMCs from rhesus macaques immunized with circRNA^RBD^ vaccines via ELISPOT assay. In **f**, the data were shown as the mean ± S.E.M. (n >= 2), and each symbol represented an individual rhesus macaque. **g**. Measuring the viral loads (N gene) and subgenome RNA loads (E gene) in the lung tissues of challenged rhesus macaques. In **g**, the data were shown as the mean ± S.E.M. (n = 4), and each symbol represented an individual rhesus macaque. **h**, The HE staining of the pathological sections using the lung tissues from immunized rhesus macaques at 7 days post SARS-CoV-2 virus challenges. **i**, The pathological score of pneumonia based on the lung tissues from immunized rhesus macaques at 7 days post SARS-CoV-2 challenges. The data were shown as the mean ± S.E.M. (n = 4), and each symbol represented an individual rhesus macaque. **j**, Correlation of B cell response, T cell response and pathological score in each immunized rhesus macaque. B cell responses were shown by the neutralizing antibody production as value of NT50 against authentic SARS-CoV-2 virus. T cell responses were shown as spots per 10^6^ PBMCs detected in IFN-γ and IL-2 ELISpot assay. Pathological scores were the same as in **i**.

We then performed the cross-neutralizing assay using the plasma samples from the immunized rhesus macaques. Both the pseudotyped and authentic SARS-CoV-2 neutralization assay showed that the circRNA^RBD^ vaccine-immunized rhesus macaque plasma could effectively inhibit the corresponding native strain, while Alpha, Delta and Beta variants could also be inhibited, but with reduced activity, especially for the Beta variant (**Fig. 5d, e**).

The peripheral blood mononuclear cells (PBMCs) collected on the day before challenge with SARS-CoV-2. The RBD-specific T cell responses elicited by circRNA^RBD^ vaccines or controls in rhesus macaques were measured using the PBMCs stimulated with the corresponding RBD peptide pools (**Supplementary Table 2**). ELISpot assay showed evident IFN-γ and IL-2 responses, but nearly undetectable IL-4 in circRNA^RBD^-immunized rhesus macaques (**Fig. 5f**), indicating a Th1-biased T cell immune response.

### circRNA vaccine protected the rhesus macaques against SARS-CoV-2 infection

Five weeks post the boost dose, the immunized rhesus macaques were challenged with 1×10^6^ plaque forming units of SARS-CoV-2 native strain via intranasal and intratracheal routes, as described previously^33^. The challenged rhesus macaques were euthanized at 7 dpi, and the lung tissues were underwent viral load and histopathological assays. The RT-qPCR assay using primers targeting SARS-CoV-2 genomic RNA (N protein region) indicated that the rhesus macaques immunized with 100 µg or 500 µg of circRNA^RBD^ vaccine were well protected as the viral genomic RNA copies were reduced nearly 1000-fold compared to the control groups (**Fig. 5g**). To detect the actively replicative viral loads, we performed qPCR using primers targeting SARS-CoV-2 sub-genomic RNA (E protein region), and found that circRNA^RBD^-immunized rhesus macaques of all three doses had nearly no detectable viral sub-genomic RNA in the lung tissues (**Fig. 5g**).

Further histopathological examination demonstrated that circRNA^RBD^-immunized rhesus macaques of all doses were well protected because only very mild pneumonia was observed, especially in the two high-dose groups (**Fig. 5h**). In contrast, severe pneumonia symptoms were observed in the lungs of two control groups, as exemplified by the local pulmonary septal thickening, moderate hemorrhage in the pulmonary septals, a large number of scattered dust cells, and massive inflammation cells infiltration (**Fig. 5h**). Pathological score further confirmed that circRNA^RBD^ immunization significantly protected the rhesus macaques from SARS-CoV-2 infection (**Fig. 5i**), likely resulting from a synergy between the humoral immune responses and T cell responses elicited by vaccination (**Fig. 5j**).

### circRNA vaccine did not cause clinical signs of illness in rhesus macaques

To further evaluate the immunogenicity and safety of circRNA vaccines in NHP, physiological and biochemical indicators were monitored, including adverse effects, cytokines indicating innate immune activation, body weight, body temperature, and blood routine examination. No severe clinical adverse effects were observed following the priming dose or the 2^nd^ boost. ELISA results showed that circRNA^RBD^ vaccines induced high levels of IL-6 and MCP-1 (**Extended Data Fig. 12a, b**), while the TNF-α, IL-1β, and IFN-α were nearly undetectable (**Extended Data Fig. 12c-e**). Body temperatures of both immunized rhesus macaques and controls were within the normal range, which have been continuously monitored for 3 days after prime and boost (**Extended Data Fig. 12f**). None of the challenged macaques showed clinical signs of illness (**Extended Data Fig. 13a-e**). Collectively, our study provided preliminary proof of safety for the circRNA vaccination in NHPs.

### Expression of SARS-CoV-2 neutralizing antibodies via circRNA platform

Besides vaccine, circRNA could be re-purposed for therapeutics when used to express other proteins or peptides, such as enzymes for rare diseases and antibodies for infectious diseases or cancer. Here, we attempted to test the therapeutic potential of circRNAs by expressing the SARS-CoV-2 neutralizing antibodies. It has been reported that SARS-CoV-2 neutralizing nanobodies or hACE2 decoys could inhibit the SARS-CoV-2 infection^73–75^. This prompted us to leverage the circRNA platform to express SARS-CoV-2 neutralizing nanobodies, including nAB1, nAB1-Tri, nAB2, nAB2-Tri, nAB3, and nAB3-Tri^73,74^, together with hACE2 decoys^75^ (**Extended Data Fig. 14a**). Pseudovirus neutralization assay showed that supernatants of HEK293T cells transfected with circRNA^nAB^ or circRNA^hACE2 decoys^ could effectively inhibit wild SARS-CoV-2 S-protein based pseudovirus infection (**Extended Data Fig. 14b**).

Next, we tested neutralizing antibodies against the recently emerged SARS-CoV-2 variants, including Alpha variant and Beta variant, by pseudovirus assays. Consistent with other reports^73,74^, the supernatants of circRNA^nAB1-Tri^ and circRNA^nAB3-Tri^ transfected cells effectively blocked Alpha and D614G pseudovirus infection (**Extended Data Fig. 14c**). However, both nanobodies showed markedly decreased neutralizing activity against Beta variant (**Extended Data Fig. 14c**). The hACE2 decoys showed no inhibition activity against Alpha and Beta variants (**Extended Data Fig. 14c**).

## Discussion

COVID-19 is still a fast-growing global health crisis with circulating SAS-CoV-2 variants evading immunity from prior vaccination or viral infection, especially with the emerging Delta and Omicron variants of concern^76–79^. The Omicron variant has been reported to escape most of SARS-CoV-2 neutralizing antibodies and the sera from vaccinees or convalescent patients^4–8^. Our study established a circular RNA vaccination strategy to elicit effective neutralizing antibodies and T cell immune responses against SARS-CoV-2 and its emerging variants (**Fig. 4** and **Fig. 5**).

As reported, most effective neutralizing antibodies recognize the RBD region of Spike protein^73,74,80–83^, and targeting RBD may induce fewer non-neutralizing antibodies^29–32,84^. Given that RBD trimers bind hACE2 better than monomeric counterparts^54^ and have been shown to enhance humoral immune response^32,54,85^, we chose to express RBD trimers via circRNA as the immunogen. The circRNA-encoded RBD trimmers were functional (**Fig. 1d**) and indeed induced sustained high-level neutralizing antibodies and specific T cell immune responses against SARS-CoV-2 and variants in both mice and rhesus macaques (**Fig. 2**, **Fig. 5** and **Extended Data Fig. 10**).

The mRNA vaccines based on the full-length Spike protein (mRNA-1273 and BNT162b2)^27,28,33^ or RBD elicit neutralizing antibodies and T cells immune responses^29–32,84^. In comparison with the mRNA vaccine, circRNA vaccine elicited longer-lasting and higher level of immunogens (**Fig. 3a-d** and **Extended Data Fig. 7**), leading to stronger Th1-biased T cell immune responses (**Fig. 3e-g**, **Extended Data Fig. 8**), suggesting that circular RNA vaccine might have a better chance to avoid the vaccine-associated enhanced respiratory disease (VAERD)^27,32,64^. Moreover, circRNA^RBD-Delta^ vaccine induced higher proportion of neutralizing antibodies against both Delta and Omicron variants than mRNA^RBD-Delta^ vaccine (**Fig. 3h-k** and **Extended Data Fig. 11**), suggesting that the circRNA vaccine platform might have superiority to cope with the potential ADE effects and emerging variants^65,66^.

It’s worthy of note that the mRNA vaccine we used for the comparison study is different from the two widely inoculated vaccines, mRNA-1273 and BNT162b2, both of which encode the full-length Spike antigens and were produced by different manufacturing process^27,28,33^.

Recently, Omicron variant rose rapidly, accounting for ∼30% of total COVID-19 cases worldwide as of Dec. 28, 2021 (GISAID). Omicron has been reported to escape most neutralizing antibodies and the inhibition from convalescent sera^4–8^. The fact that circRNA^RBD-Omicron^ vaccine we developed could not cross-protect Delta variant (**Fig. 4d-f**) suggests that the immunogenicity of Omicron RBD was less effective and largely different from Delta RBD^3^.

A recent preprint reported that vaccinees who received two doses of SARS-CoV-2 vaccine exhibited enhanced neutralizing antibodies against Delta variant after infected by Omicron, implying that Omicron vaccine might provide broad-spectrum protection against other variants^86^. Our result argues against such possibility because our Omicron-specific vaccine failed to cross protect Delta variant (**Fig. 4d-f**), or boost two-dose of Delta vaccine (**Fig. 4h, i**). On the contrary, circRNA^RBD-Delta^ vaccine appeared to produce antigens possessing high immunogenicity and consequently elicit high level of neutralizing antibodies against Delta (**Fig. 2**). Our Delta-specific vaccination could cross protect all other variants including Omicron (**Fig. 2** and **Fig. 4a-c**), and it could also be used as an effective booster following two-dose original SARS-CoV-2 vaccines (**Fig. 4k, l**). It is hopeful but remains to be further tested whether circRNA^RBD-Delta^ vaccine could be applied as an effective booster for current major vaccines, which are exclusively based on the original SARS-CoV-2 strain.

It has been a concerning issue whether the circRNAs produced via *in vitro* transcription induce strong innate immune responses^38,39,87^. Our vaccination study in rhesus macaques demonstrated that circRNA vaccine could indeed provide safe protection against SARS-CoV-2 infection without causing severe clinical signs of illness (**Fig. 5**, **Extended Data Fig. 12a-f** and **Extended Data Fig. 13a-e**).

In this study, we also tested the therapeutic potential of circRNAs that encode SARS-CoV-2-specific neutralizing nanobodies^73,74,80–83^ or hACE2 decoys^88,89^, which could effectively neutralize the SARS-CoV-2 pseudovirus **(Extended Data Fig. 14a-c)**. Beyond viral receptors, this circRNA expression platform hold the potential to become therapeutic drugs, encoding therapeutic antibodies *in vivo,* such as anti-PD1/PD-L1 antibodies^90,91^. Unlike antibodies and protein drugs, circRNAs encode therapeutic antibodies in the cytoplasm, allowing them to target intracellular targets like TP53^92^ and KRAS^93^, bypassing the cytomembrane barrier.

Collectively, circular RNA holds the potential to become an effective and safe vaccine platform against viral infection, including SARS-CoV-2 emerging variants, as well as a therapeutic platform, owing to its specific properties.

---

**Extended Data Fig. 1.**
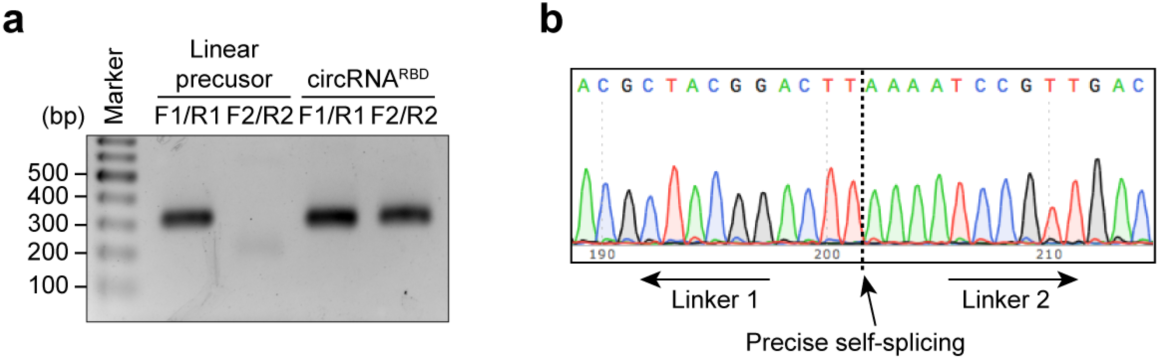
PCR analysis to verify the precise circularization of circRNA. **a**, The agarose gel electrophoresis result of PCR analysis. Linear RNA precusor and circRNA^RBD^ were reverse transcription to cDNA, followed by PCR amplification with specific primers shown in **Fig 1a**. **b**, Sanger sequence result of the PCR products in (**a**).

**Extended Data Fig. 2.**
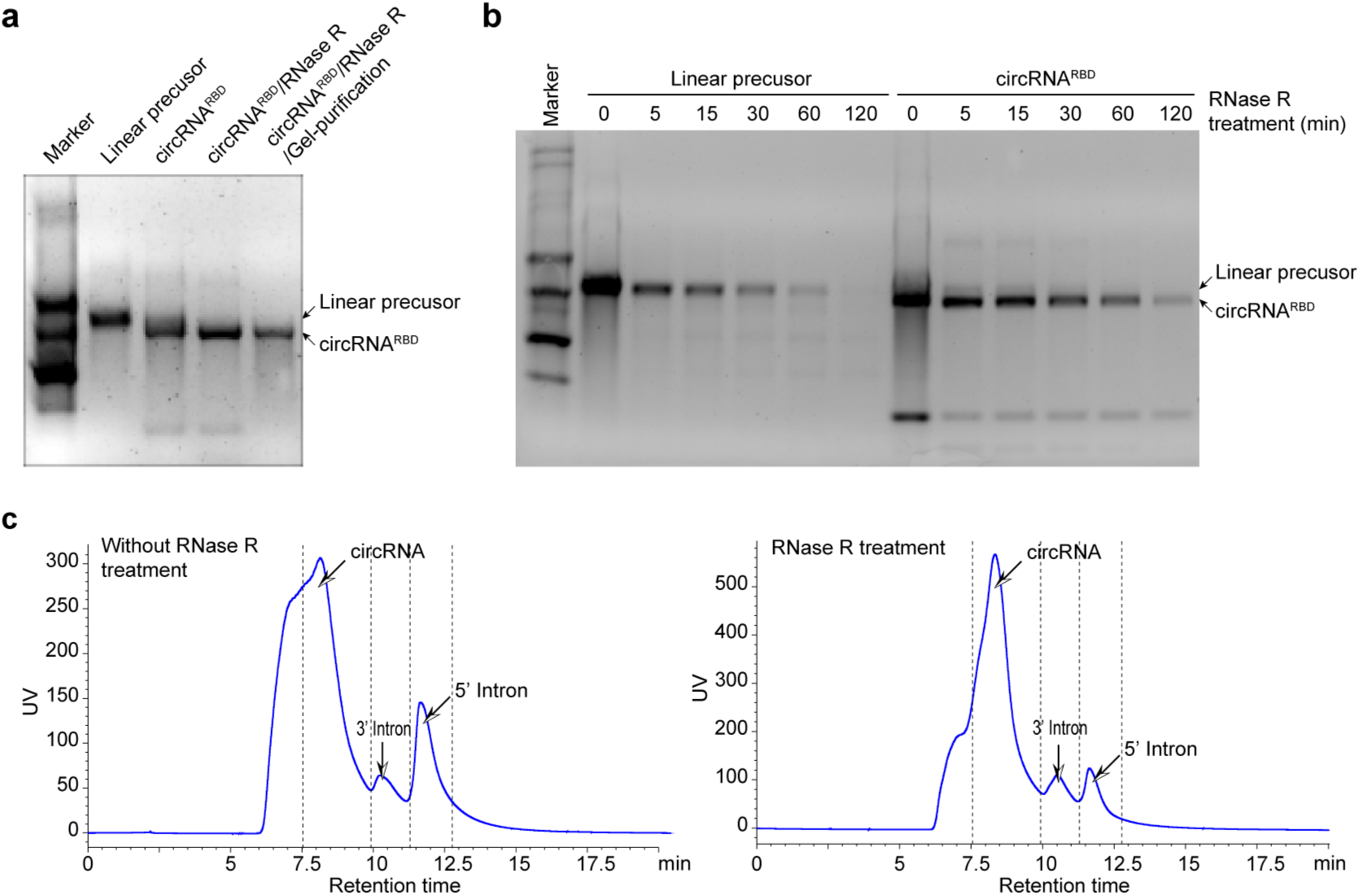
Agarose gel electrophoresis and HPLC purification of circRNA^RBD^. **a**, The agarose gel electrophoresis result of linear RNA precusor and circRNA^RBD^ with different treatments. **b**, The agarose gel electrophoresis result of circRNA^RBD^ and linear RNA precusor digested by RNase R for various time from 5 min to 120 min. **c**, HPLC chromatogram of circRNA^RBD^ without RNase R treatment (left) and circRNA^RBD^ treated by RNase R (right).

**Extended Data Fig. 3.**
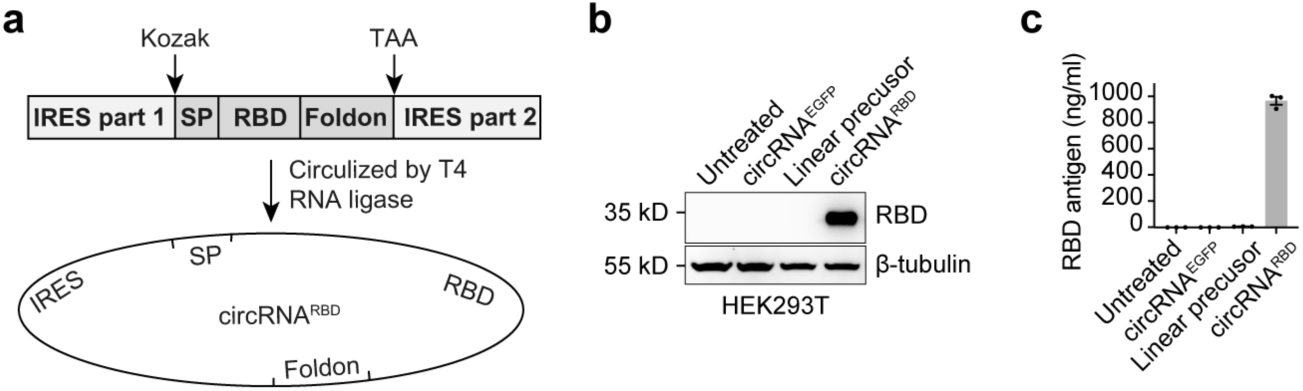
Expression of SARS-CoV-2 RBD antigens with circular RNAs circulized by T4 RNA ligase. **a**, Schematic diagram of circRNA^RBD^ circularization by the T4 RNA ligase. SP, signal peptide sequence of human tPA protein. Foldon, the trimerization domain from bacteriophage T4 fibritin protein. RBD, the receptor binding domain of SARS-CoV-2 Spike protein. **b**, Western Blot analysis showing the expression level of RBD antigens in the supernatant of HEK293T cells transfected with circRNA^RBD^ circulized by the T4 RNA ligase. The circRNA^EGFP^ and linear RNA precursor were used as controls. **c**, Measuring the concentration of RBD antigens in the supernatant with quantitative ELISA assay. In **c**, data were shown as the mean ± S.E.M. (n = 3).

**Extended Data Fig. 4.**
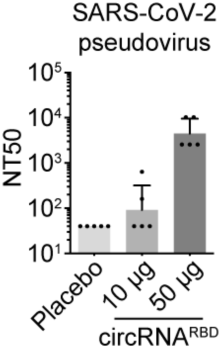
Measuring the NT50 value of sera from mice immunized with cirRNA^RBD^ vaccine. The NT50 value was calculated using lentivirus-based SARS-CoV-2 pseudovirus. Data were shown as the geometric mean ± geometric S.D., and each symbol represented an individual mouse.

**Extended Data Fig. 5.**
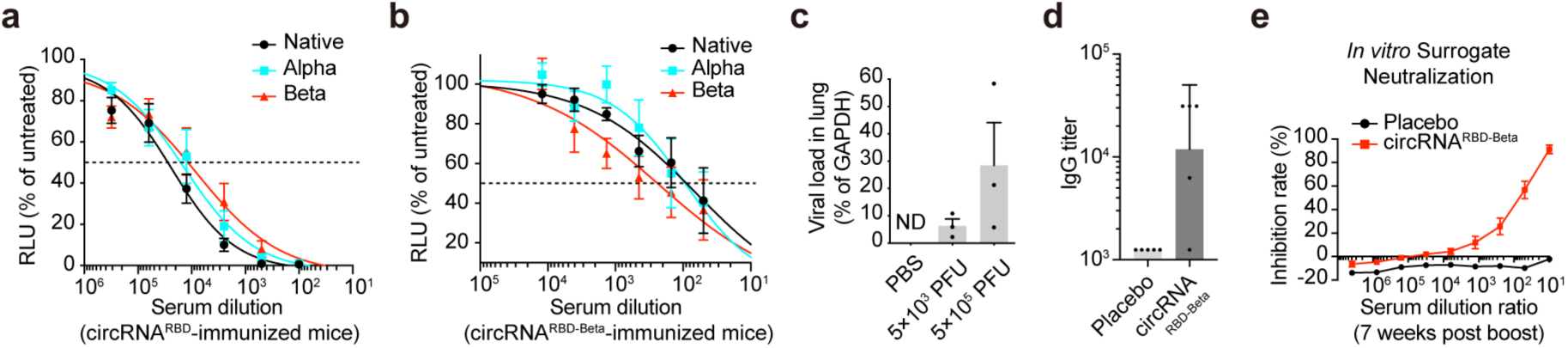
SARS-CoV-2 circRNA^RBD-Beta^ vaccine-elicited antibodies showed preferential neutralizing activity against Beta variant. **a** and **b**, Sigmoidal curve diagram of neutralization rate of VSV-based D614G, Alpha or Beta pseudovirus with the sera of mice immunized with circRNA^RBD^ or circRNA^RBD-Beta^ vaccines. The data were shown as the mean ± S.E.M. (n = 4). Neutralization assay of the serum samples were collected at 1 week post boost. The data were shown as the mean ± S.E.M. (n = 5). **c**, Measuring the viral loads in the lung tissues of BALB/c mice infected by authentic SARS-CoV-2 Beta virus. The SARS-CoV-2 RNA copies were normalized with *GAPDH*. The data were shown as the mean ± S.E.M. (n = 3). **d**, Measuring the SARS-CoV-2 RBD-Beta-specific IgG endpoint GMTs with ELISA. In **d**, data were shown as the geometric mean ± geometric S.D., and each symbol represented an individual mouse. **e**, Sigmoidal curve diagram of the inhibition rate by sera from immunized mice with surrogate virus neutralization assay. In **e**, data were shown as the mean ± S.E.M. (n = 5). In **d** and **e**, the sera from circRNA^RBD-Beta^ (50 µg) immunized mice were collected at 3 days before challenge with authentic SARS-CoV-2 Beta strain.

**Extended Data Fig. 6.**
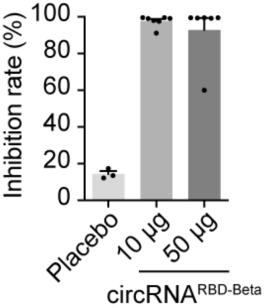
Measuring the neutralizing activity of sera from circRNA^RBD-Beta^-immunized mice with surrogate SARS-CoV-2 neutralization assay. Instead of the lab-prepared LNP, the commercial LNP (Precision Nanosystems) was used to encapsulate the circRNA^RBD-Beta^ vaccine, and then the mice were immunized i.m. with 10 µg or 50 µg of new LNP-circRNA^RBD-Beta^ vaccine at a two-week interval. The sera of immunized mice were collected at 2 weeks post the second dose. The data were shown as the mean ± S.E.M. Each symbol represented an individual mouse.

**Extended Data Fig. 7.**
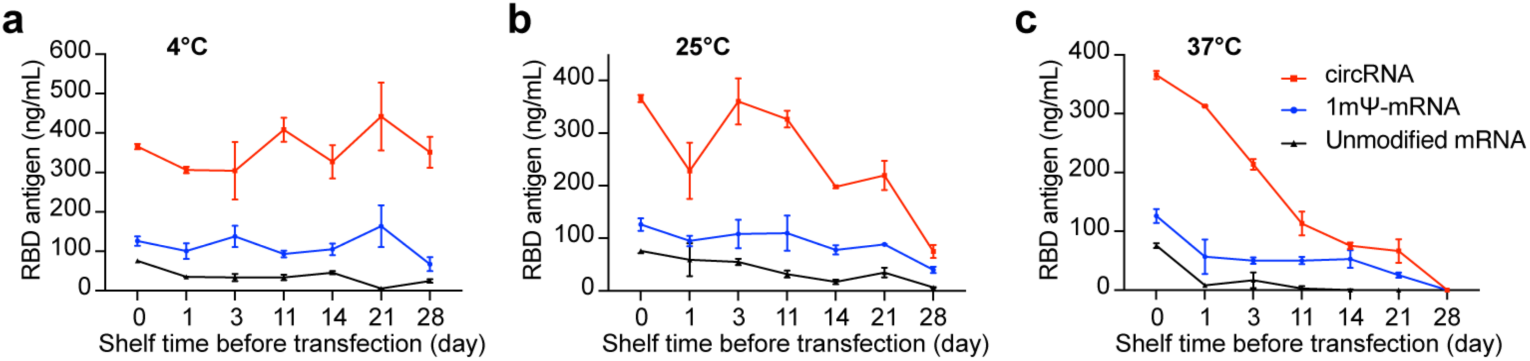
Measuring the expression level of RBD-Delta antigens in the supernatant of HEK293T cells at different storage condition. **a**-**c**, Using quantitative ELISA to measure the expression level of RBD-Delta antigens in the supernatant of HEK293T cells transfected with LNP-circRNA^RBD-Delta^, LNP-1mΨ mRNA^RBD-Delta^ and LNP-non-modified-mRNA^RBD-Delta^, stored at 4°C (**a**), 25°C (**b**) or 37°C (**c**). The LNP-RNA were stored at different temperature and transfected at different time points. The data were shown as the mean ± S.E.M. (n = 3).

**Extended Data Fig. 8.**
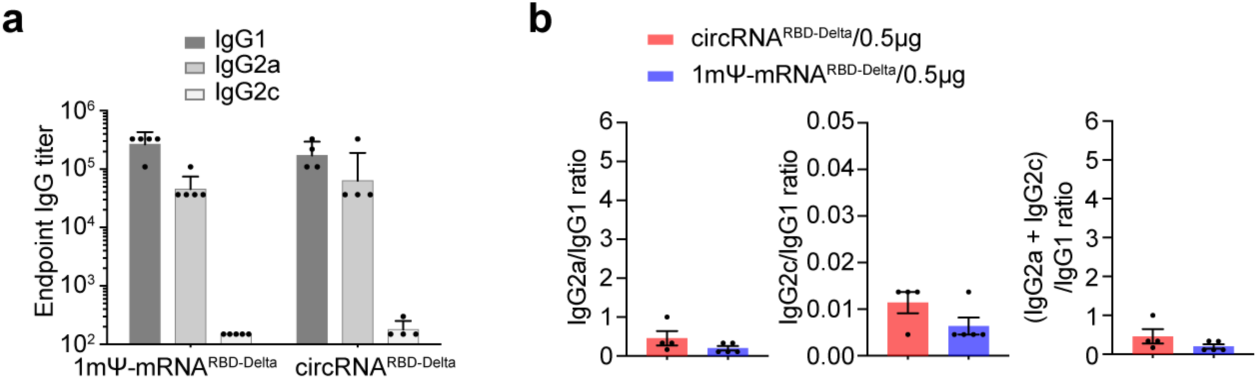
The circRNA vaccine elicited stronger Th1-biased T cell immune responses than 1mΨ-mRNA vaccine. **a**, Measuring RBD-Delta-specific IgG1/IgG2a/IgG2c endpoint GMTs elicited by 0.5 µg of circRNA^RBD-Delta^ vaccine or 1mΨ-mRNA^RBD-Delta^ vaccine in mice. In **a**, data were shown as the geometric mean ± geometric S.D., and each symbol represented an individual mouse. **b**, Measuring the specific IgG2a/IgG1, IgG2c/IgG1 and (IgG2a + IgG2c)/IgG1 ratio of serum from mice immunized with 0.5 µg of circRNA^RBD-Delta^ or 1mΨ mRNA^RBD-Delta^. In **b**, data were shown as the mean ± S.E.M., and each symbol represented an individual mouse.

**Extended Data Fig. 9.**
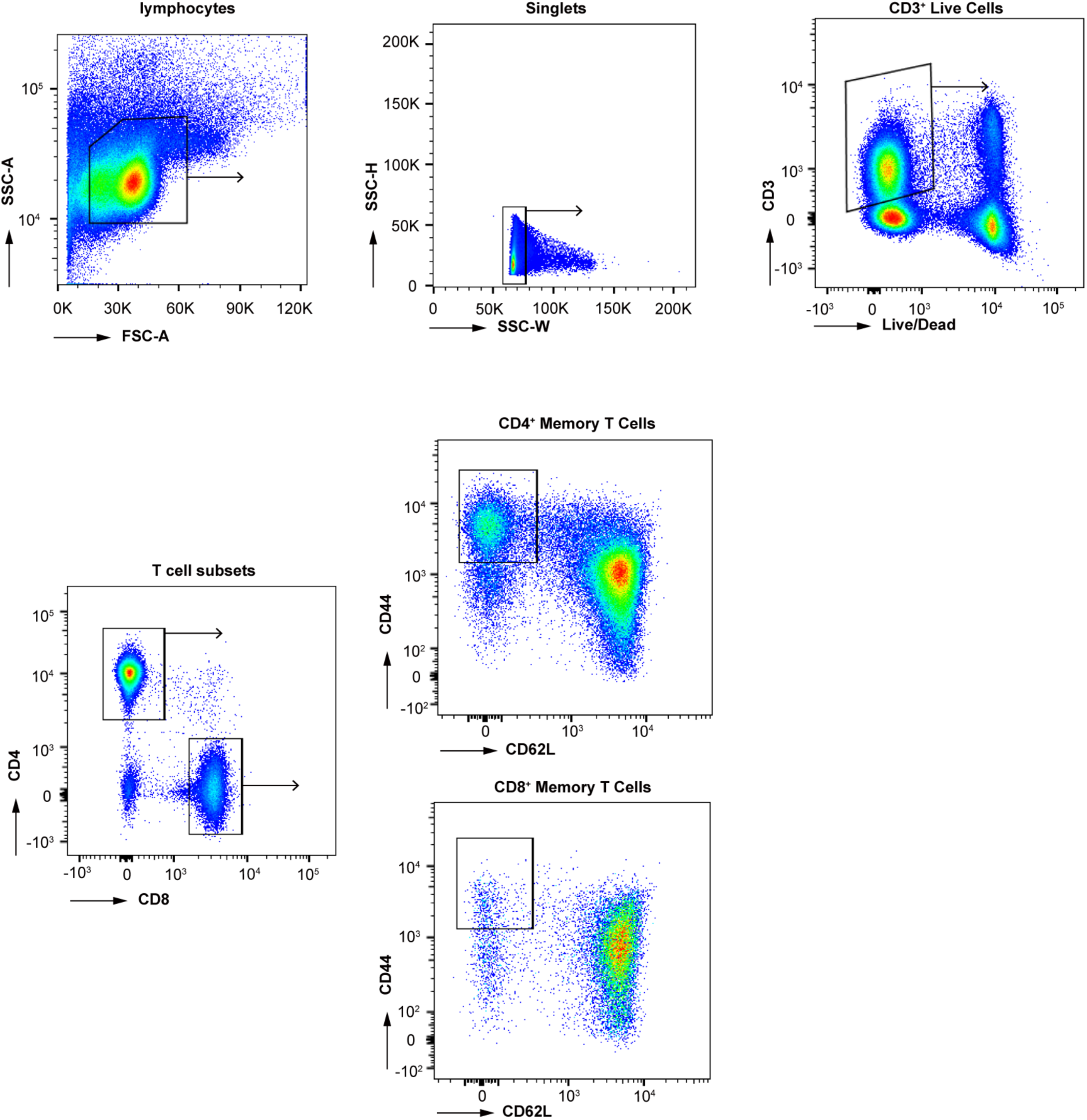
Flow panel and gating strategy to quantify SARS-CoV-2-RBD-specific T cells in mice. The plots showed the gating strategy of single and viable T cells in spleenocytes. CD4^+^ or CD8^+^ Tem cells (CD44^+^CD62L^-^) were further analyzed for detecting the expression of cytokines stimulated by corresponding RBD-Delta peptide pools.

**Extended Data Fig. 10.**
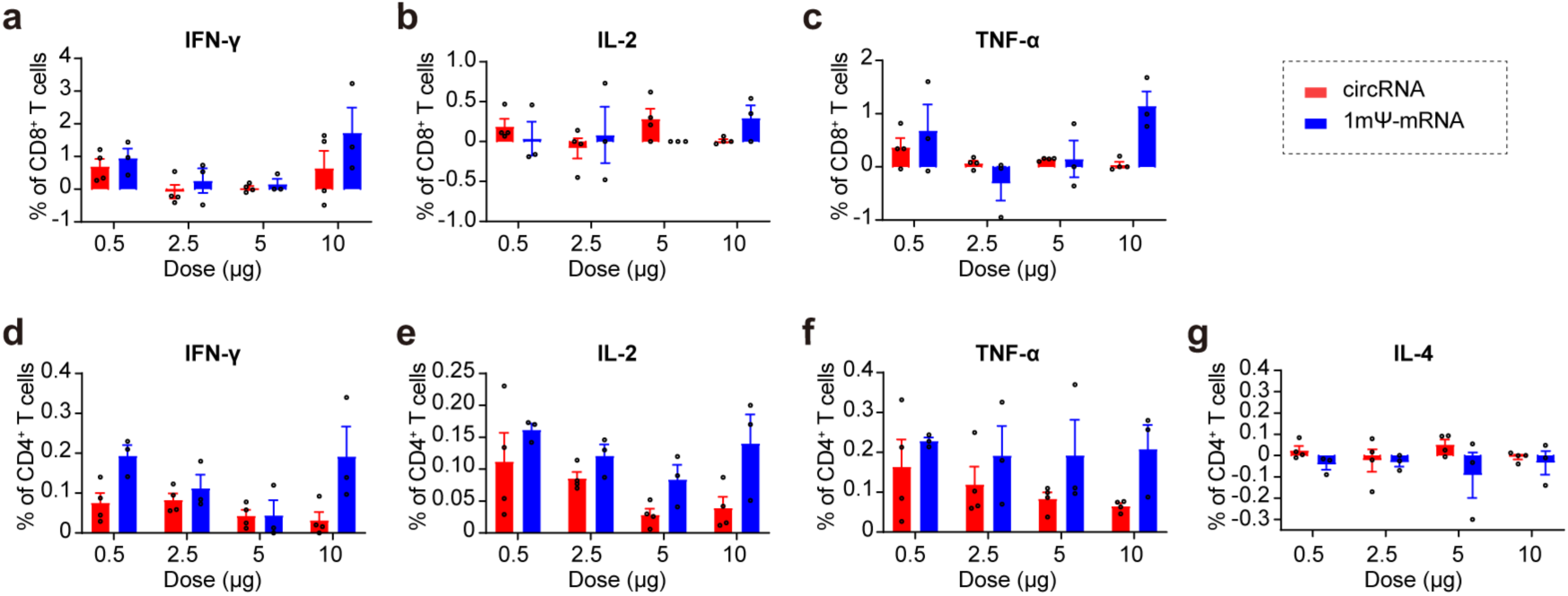
T cell immune responses elicited by SARS-CoV-2 circRNA^RBD-Delta^ or mRNA^RBD-Delta^ vaccines in mice. **a**-**c**, The FACS analysis results showing the percentages of CD8^+^ Tem cells secreting IFN-γ (**a**), IL-2 (**b**), or TNF-α (**c**) after stimulated by RBD-Delta peptide pools, respectively. **d**-**g**, The FACS analysis results showing the percentages of CD4^+^ Tem cells secreting IFN-γ (**d**), IL-2 (**e**), TNF-α (**f**) or IL-4 (**g**) after stimulated by RBD-Delta peptide pools, respectively. In **a-g**, the data were presented as the mean ± S.E.M. (n = 3 or 4), and each symbol represented an individual mouse.

**Extended Data Fig. 11.**
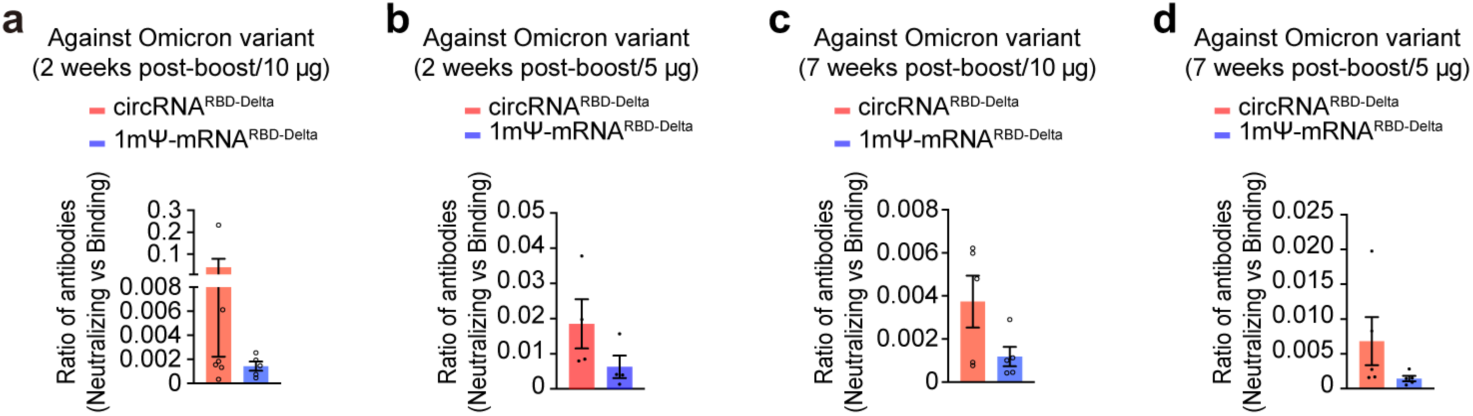
The circRNA^RBD-Delta^ vaccine elicited high-level of neutralizing antibodies against Omicron variant. **a** and **b**, Measuring the ratio of (Neutralizing antibodies)/(Binding antibodies) elicited by 10 µg (**a**) or 5 µg (**b**) of circRNA^RBD-Delta^ vaccine or 1mΨ-mRNA^RBD-Delta^ vaccine with the sera collected at 2 weeks post boost. The ratio of (NT50)/(Endpoint GMT) of each mouse was calculated. **c** and **d**, Measuring the ratio of (Neutralizing antibodies)/(Binding antibodies) elicited by 10 µg (**c**) or 5 µg (**d**) of circRNA^RBD-Delta^ vaccine or 1mΨ-mRNA^RBD-Delta^ vaccine with the sera collected at 7 weeks post boost. The ratio of (NT50)/(Endpoint GMT) of each mouse was calculated. In **a**-**d**, data were presented as the mean ± S.E.M., and each symbol represented an individual mouse.

**Extended Data Fig. 12.**
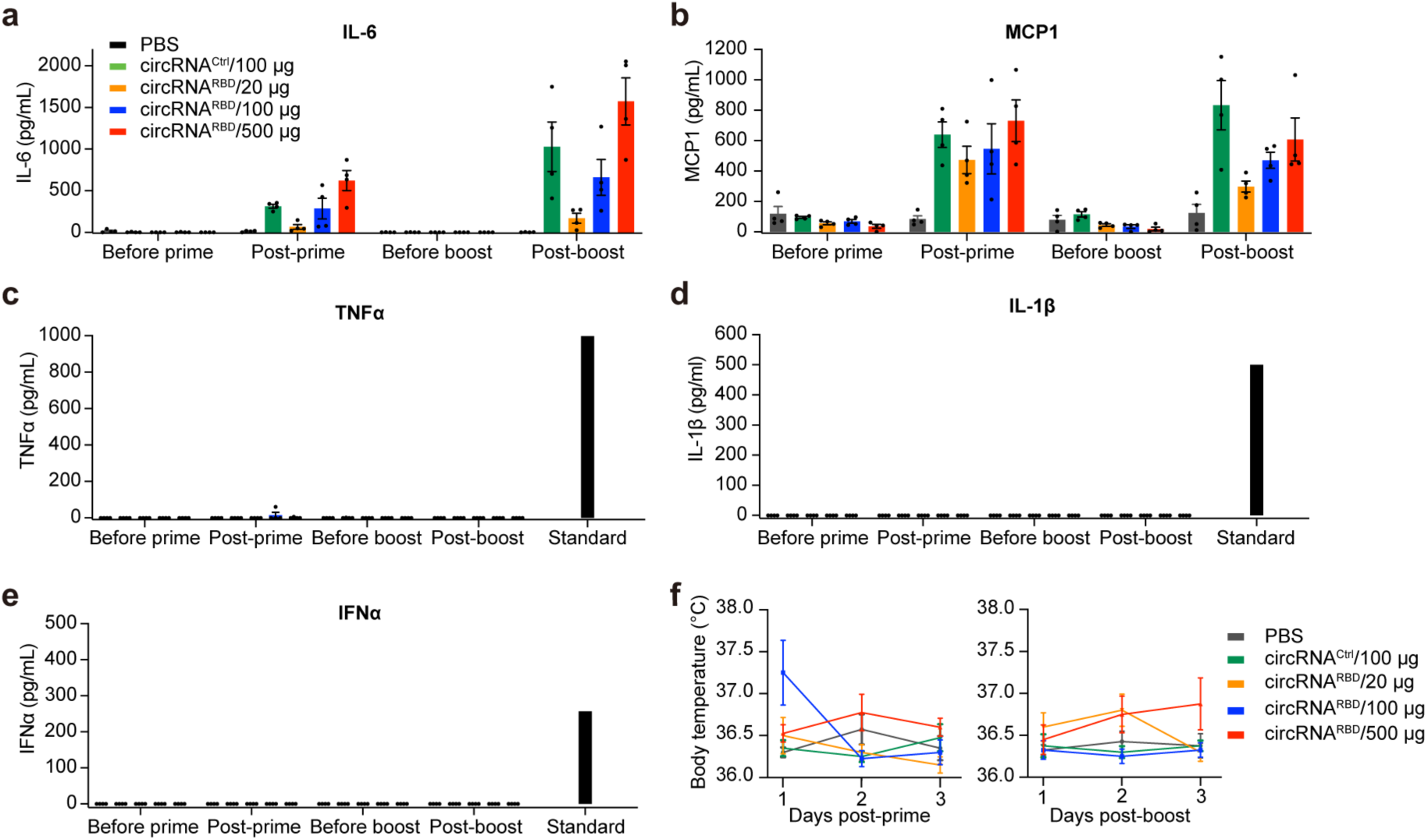
CircRNA vaccine caused no obvious clinical signs of illness in rhesus macaques. **a**-**e**, Measuring the IL-6 (**a**), MCP1 (**b**), TNF-α (**c**), IL-1β (**d**) and IFN-α (**e**) level in the plasma of immunized rhesus macaques. **f**, Monitoring the body temperature of rhesus macaques. The body temperature is monitored within three days after the prime and boost dose. In **a**-**f**, data were shown as the mean ± S.E.M. (n = 4).

**Extended Data Fig. 13.**
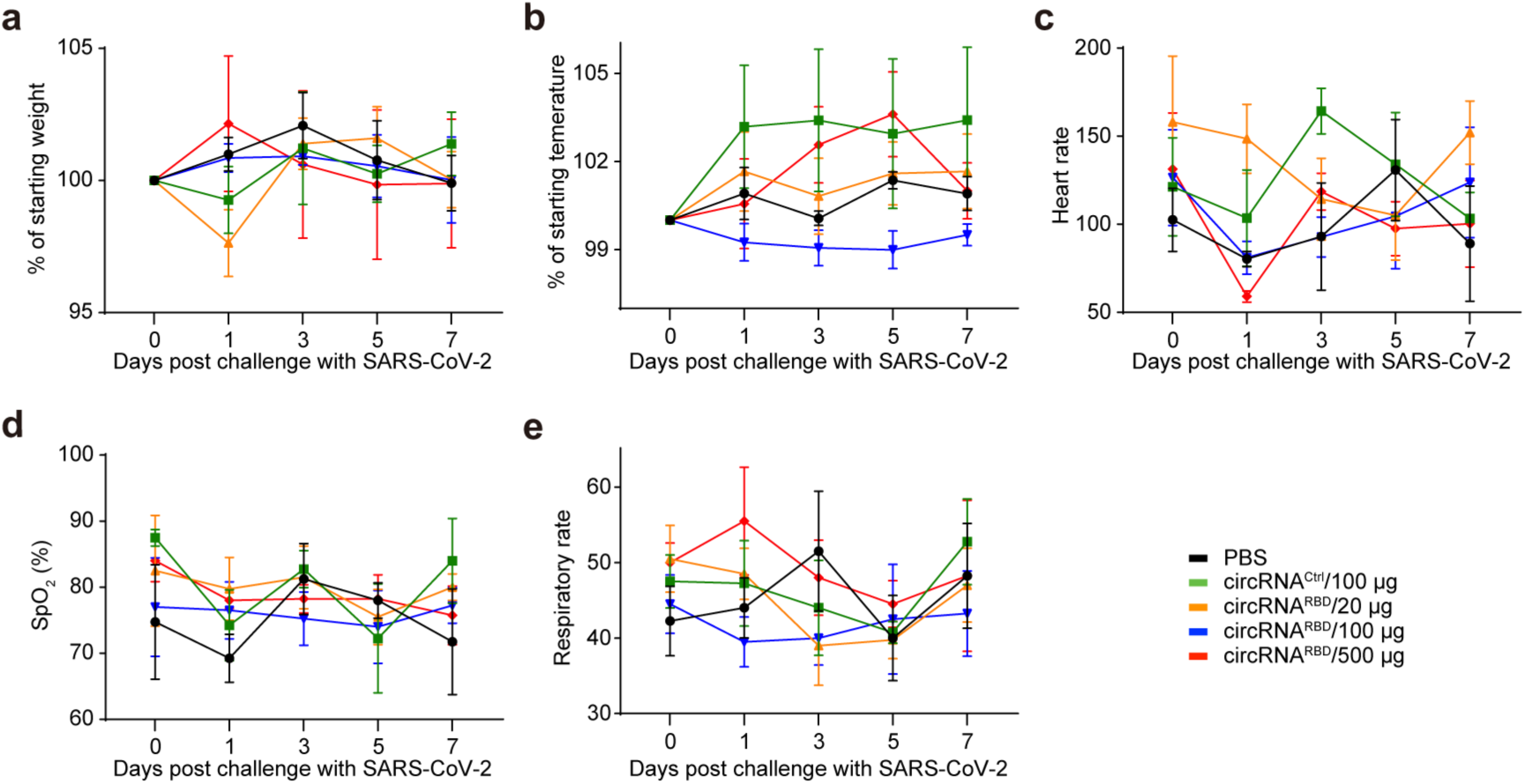
Vital clinical signs of circRNA^RBD^-immunized rhesus macaques after challenge with SARS-CoV-2. **a-e**, The body weight (**a**), temperature (**b**), heart rate (**c**), oxygen saturation (**d**), respiratory rate (**e**) were monitored after challenge with SARS-CoV-2 virus. In **a**-**e**, data were shown as the mean ± S.E.M. (n = 4).

**Extended Data Fig. 14.**
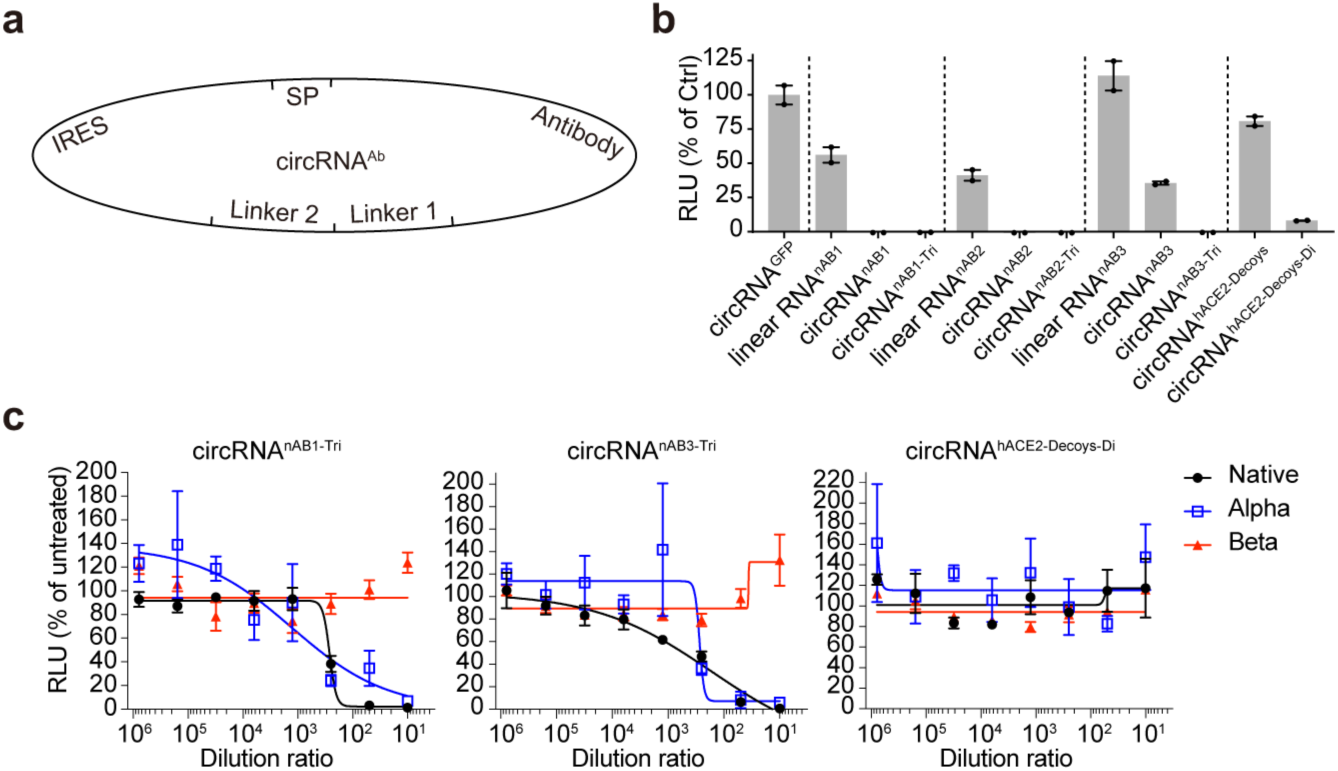
Expression of SARS-CoV-2 neutralizing nanobodies or hACE2 decoys via circRNA platform. **a**, Schematic diagram of circRNA^nAB^ or circRNA^hACE2 decoys^ circularization by the Group I ribozyme autocatalysis. **b**, Lentivirial-based pseudovirus neutralization assay with the supernatant from cells transfected with circRNA encoding neutralizing nanobodies nAB1, nAB1-Tri, nAB2, nAB2-Tri, nAB3 and nAB3-Tri or ACE2 decoys. The nAB1-Tri, nAB2-Tri and nAB3-Tri represented the trimmer of nAB1, nAB2 and nAB3, respectively. The luciferase value was normalized to the circRNA^EGFP^ control. In **b**, data were shown as the mean ± S.E.M. (n = 2). **c**, Sigmoidal curve diagram of neutralization rate of VSV-based SARS-CoV-2 D614G, Alpha or Beta pseudovirus using the supernatant of cells transfected with neutralizing nanobodies nAB1-Tri, nAB3-Tri or ACE2 decoys encoded by the corresponding circRNAs. In **c**, data were shown as the mean ± S.E.M. (n = 3).

**Supplementary Table 1.**
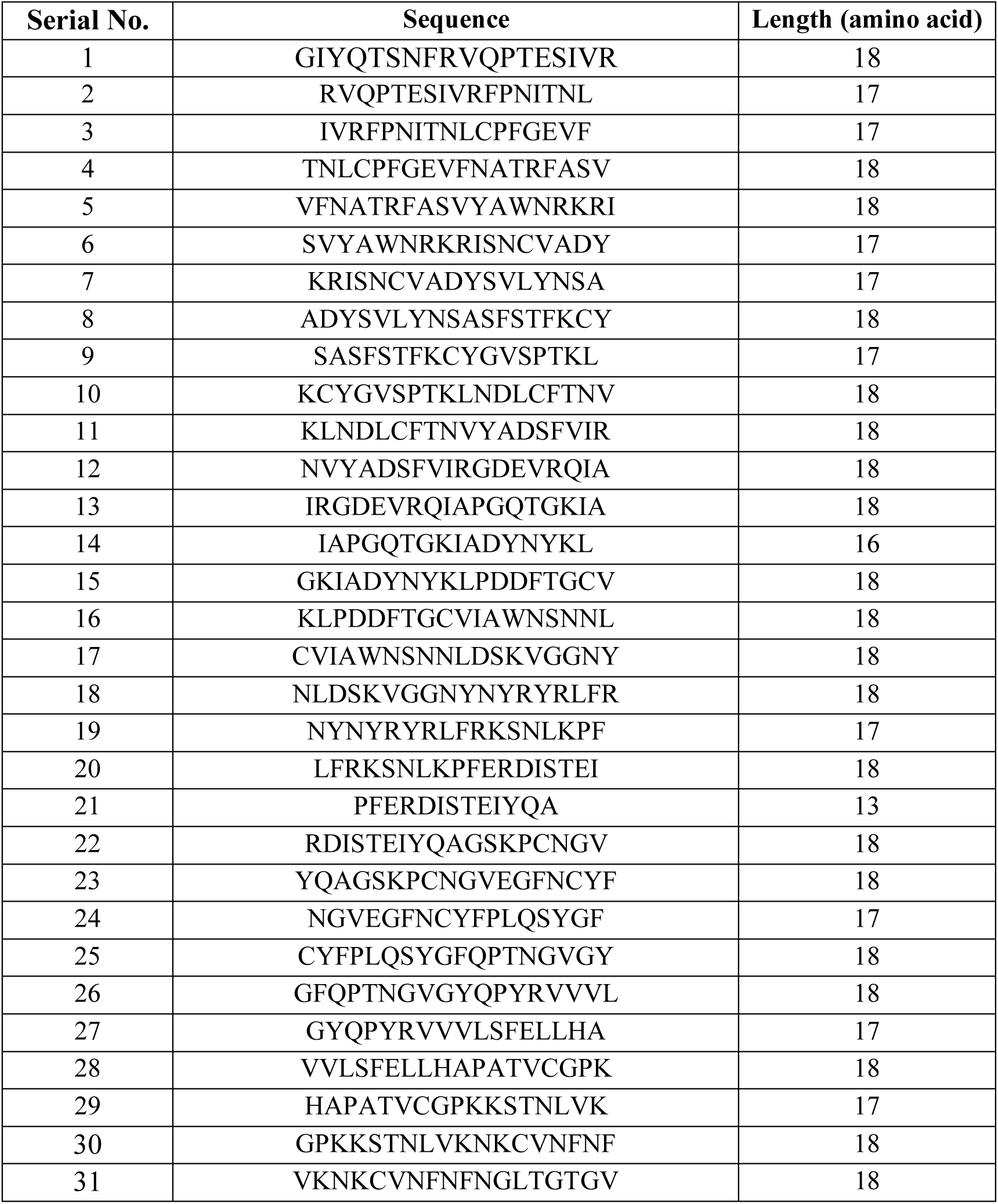
The peptide sequences of RBD-Delta antigen.

**Supplementary Table 2.**
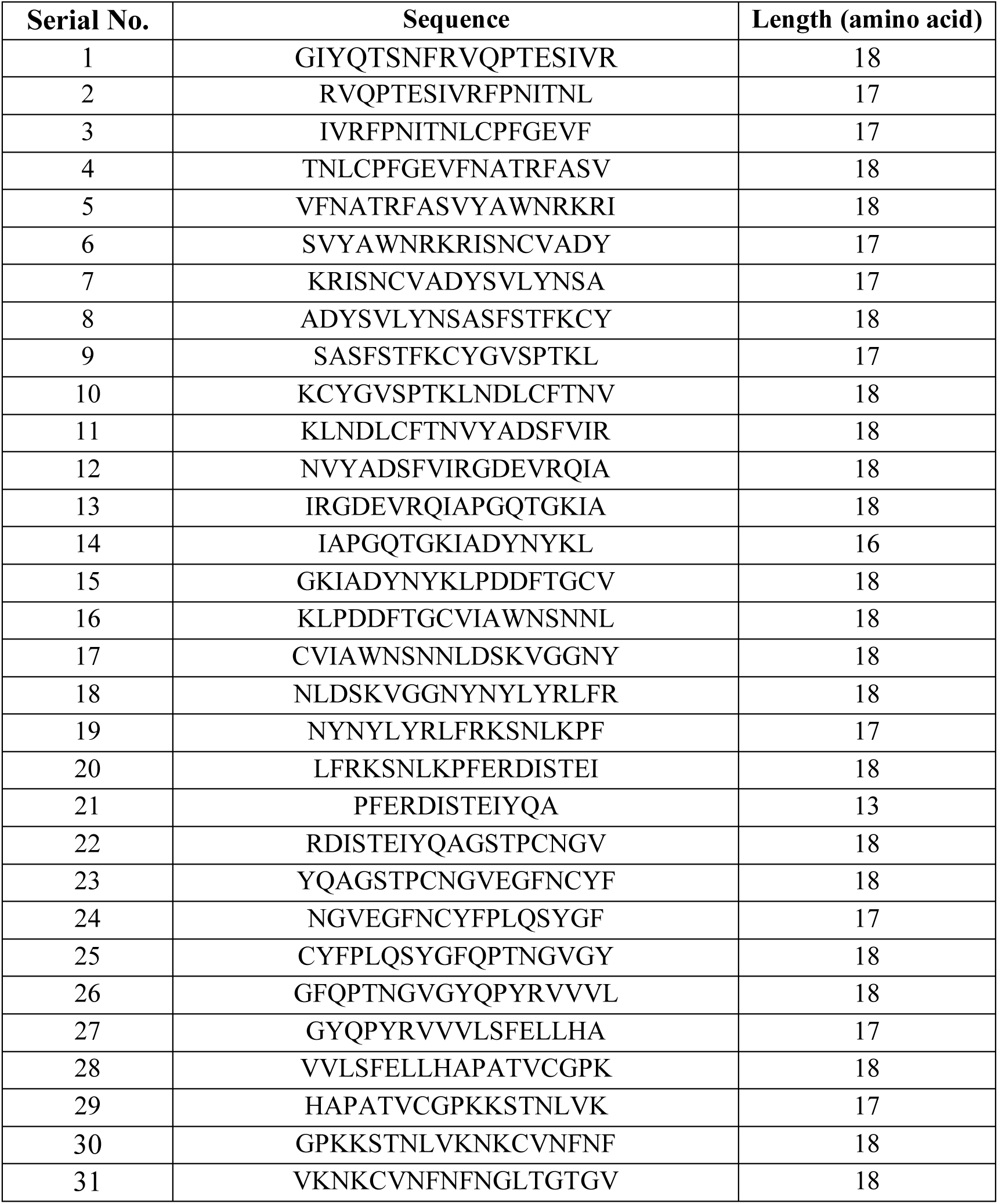
The peptide sequences of RBD antigen.

## Acknowledgements

We acknowledge C. Dong and X. Wang (Tsinghua University) for providing fluorescent-conjugated antibodies. We thank J. Xiao (Peking University) for providing purified SARS-CoV-2 Spike proteins. We thank J. Yan (Institute of Microbiology, Chinese Academy of Sciences) for providing RBD peptide pools. We thank the Laboratory Animal Centre of Peking University for the feeding of mice. We thank the HPLC Core at the National Center for Protein Sciences at Peking University (Beijing), particularly H. Li and G. Li for technical help. We thank the flow cytometry Core at the National Center for Protein Sciences at Peking University (Beijing), particularly H. Lv, Y. Guo, H. Yang, F. Wang for technical help. This project was supported by funds from National Key R&D Program of China (2020YFA0707800 to W.W., 2020YFA0707600 to Z.Z.); Beijing Municipal Science & Technology Commission (Z181100001318009); the National Science Foundation of China (31930016); Beijing Advanced Innovation Center for Genomics at Peking University and the Peking-Tsinghua Center for Life Sciences (to W.W.); the National Science Foundation of China (31870893); the National Major Science & Technology Project for Control and Prevention of Major Infectious Diseases in China (2018ZX10301401 to Z.Z.) and the Fellowship of China National Postdoctoral Program for Innovative Talents (BX20200010, to L.Q.).

## Author contributions

W.W. conceived and supervised this project. W.W., L.Q., Z.Y. and Y.S. designed the experiments. L.Q., Z.Y., Y.S., L.L., F.C., Y.X., Z.W., H.T., A.Y., and X.X. performed the experiments with the help from X.Z., F.T., C.W., X.D., L.G., S.L., C.Y., C.T., Y.Y., W.Y., J.W., Y.Z., Q.H., Z.Z., Y.C., Y.W., X.P., J.W., and X.S.X.. L.Q., Z.Y., Y.S. and W.W. wrote the manuscript with the help of all other authors.

## Methods

### Plasmids construction

The SARS-CoV-2 RBD antigen, EGFP, nanobody or hACE2-decoy-coding sequence was PCR amplified and cloned into the pcircRNA-EV backbone and constructed the corresponding pcircRNA plasmids for the following IVT (*in vitro* transcription).

### Cell culture

HEK293T, NIH3T3 and Huh-7 cell lines were maintained in our laboratory. The HEK293T-hACE2 cell line was ordered from Biodragon Inc. (#BDAA0039, Beijing, China). The A549-hACE2 cell line was generated in our laboratory. These mammalian cell lines were cultured in Dulbecco’s Modified Eagle Medium (Corning, 10-013-CV) with 10% fetal bovine serum (FBS) (BI), supplemented with 1% penicillin-streptomycin in 5% CO_2_ incubator at 37°C. The Huh-7 cells were cultured with the methods previously described^80^.

### circRNA transfection *in vitro*

For the circRNA transfection in HEK293T or NIH3T3 cells, 3×10^5^ cells per well were seeded in 12-well plates. 2 µg of circRNAs were transfected into the HEK293T or NIH3T3 cells, using Lipofectamine MessengerMax (Invitrogen, LMRNA003) according to the manufacturer’s instructions. At 24-48 hr post transfection, the cell lysis and supernatant were collected for the following detections.

### Quantitative determination of SARS-CoV-2 Spike RBD expression in vitro

Quantification of RBD expression in cell culture supernatants was performed with a commercial SARS-CoV-2 Spike RBD Protein ELISA kit (ABclonal, #RK04135) according to the manufacturer’s instruction. The supernatants were diluted at 1:500 rate. Final concentrations of RBD were calculated basing on the linear standard curve of absorbance at 450 nm, using 630 nm as reference. Briefly, the detection wells were pre-coated with monoclonal antibody specific for Spike RBD protein. After incubation with samples or standards at 37°C for two hours, samples unbound to immobilized antibody would be removed by washing steps. Then the RBD-specific antibodies were added to wells for one-hour incubation at 37°C. After washing, the HRP substrates and stop solution were added and the absorbance at 450 nm were measured using 630 nm as reference.

### Mouse vaccination and serum collection

The BALB/c mice were ordered from Beijing Vital River Laboratory Animal Technology Co., Ltd. All mice were bred and kept under SPF (specific pathogen-free) conditions in the Laboratory Animal Center of Peking University. The animal experiments were approved by Peking University Laboratory Animal Center (Beijing), and undertaken in accordance with the National Institute of Health Guide for Care and Use of Laboratory Animals. All the animal experiments with SARS-CoV-2 challenge were conducted under animal biosafety level 3 (ABSL3) facilities in Institute of Pathogen Biology, Chinese Academy of Medical Sciences. All the animal experiments with SARS-CoV-2 challenge were reviewed and approved by the Committee on the Ethics of Animal Experiments of Institute of Pathogen Biology, Chinese Academy of Medical Sciences.

For mouse vaccination, groups of 6-8 week-old female BLAB/c mice were intramuscularly immunized with LNP-circRNA^RBD^ or Placebo (Empty LNP) in 100 µL using a 1 mL sterile syringe, and 2 or 3 weeks later a second dose was immunized to boost the immune responses. The sera of immunized mice were collected to detect the SARS-CoV-2-specific IgG endpoint GMTs and neutralizing antibodies as described below.

### Antibody endpoint GMT measurement with ELISA

All the immunized mouse serum samples were heat-inactivated at 56°C for 30 min before use. The SARS-CoV-2-specific IgG antibody endpoint GMT was measured by ELISA. Briefly, serial 3-fold dilutions (in 1% BSA) of heat-inactivated sera, starting at 1:100, were added to the 96-well plates (100 µL/well; Costar) coated with recombinant SARS-CoV-2 Spike or RBD antigens (Sino Biological) and blocked with 1% BSA for 60 min at 37°C. Then, after three washes with wash buffer, the Horseradish peroxidase HRP-conjugated rabbit anti-mouse IgG (Sigma) diluted in 1% BSA at 1:10,000 ratio, was added to the plates and incubated at 37°C for 30 min. Then the plates were washed for 3 times with wash buffer and added with TMB substrates (100 µL/well) followed by incubation for 15-20 min. And then the ELISA stop buffer was added into the plates. Finally, the absorbance (450/630 nm) was measured with Infinite M200 (TECAN). The IgG endpoint GMTs were defined as the dilution fold, which emitted an optical density exceeding 3x background (without serum but the secondary antibody was added).

### SARS-CoV-2 Surrogate Virus Neutralization Assay

The neutralizing activity of mouse serum samples was detected by SARS-CoV-2 Surrogate Virus Neutralization Test Kit (L00847A, GenScript). Detections were performed according to manufacturer’s instruction. Serial 10-fold dilutions of heat-inactivated sera, starting at 1:10, were incubated with HRP-conjugated RBD solutions at 37°C for half an hour, and then the mixtures were added into 96-well plates pre-coated with human ACE2 (hACE2) proteins and incubated for 15 min at 37°C. After washing the TMB substrates and stop solutions were added and the absorbance (450/630 nm) was measured with Infinite M200 (TECAN). The inhibition rates of serum samples were calculated according to the following formula. The 50% neutralization geometric mean titer (NT50) was determined using four-parameter nonlinear regression in Prism 8 (GraphPad).

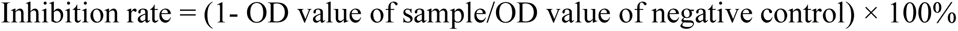

### Pseudovirus-based neutralization assay

The production of lentivirus-based SARS-CoV-2 pseudovirus and neutralization assay were performed as described previously^94^. Briefly, the SARS-CoV-2 pseudovirus were produced by co-transfecting plasmids psPAX2 (6 µg), pSpike (6 µg), and pLenti-Luc-GFP (6 µg) into HEK293T cells using X tremeGENE HP DNA Transfection Reagent (Roche) according to the manufacturer′s instructions. 48 hr post transfection, the supernatants containing pseudovirus particles were harvested and filtered through a 0.22-µm sterilized membrane for the neutralization assay as described below.

For the determination of NT50 of immunized mouse serum, the HEK293T-hACE2 cells were seeded in 96-well plates (50,000 cells/well) and incubated for approximate 24 hr until reaching over 90% confluent, preparing for pseudovirus infection. The mouse serum was 3-fold diluted, starting at 1:40, and incubated with the SARS-CoV-2 pseudovirus (MOI ≈ 0.05) at 37°C for 60 min. The DMEM medium without serum was used as the negative control group. Then the supernatant of HEK293T-hACE2 cells were removed and the mixer of serum and pseudovirus were added to each well. 36-48 hr later, the luciferase activity, which reflecting the degree of SARS-CoV-2 pseudovirus transfection, was measured using the Nano-Glo Luciferase Assay System (Promega). The NT50 was defined as the fold-dilution, which emitted an exceeding 50% inhibition of pseudovirus infection in comparison with the control group.

The neutralization assay of VSV-based pseudovirus of SARS-CoV-2 and variants was performed as described previously^80,81^. Briefly, the sera were serially diluted using complete DMEM culture media in 96-well white plates for a total of six gradients, and then the virus solution with ∼1.3×10^4^ TCID50 was added. The complete DMEM medium was used as the control group. After one hour incubation in 5% CO_2_ incubator at 37°C, the 96-well white plates were added with Huh7 cells (100 µL/well), which were adjusted to a concentration of 2×10^5^ cells/mL. After 24 h incubation in 5% CO_2_ incubator at 37°C, the culture supernatant was aspirated gently to leave 100 µL in each well, and then 100 µL of luciferase substrate (Perkinelmer, 6066769) was added to each well for the detection of luminescence using Infinite M200 (TECAN). Relative luciferase units (RLU) were normalized to corresponding DMEM control group, and the NT50 were determined by a four-parameter nonlinear regression in Prism (GraphPad).

For the neutralization assay of circRNA^nAB^ or circRNA^ACE2 decoys^, the HEK293T-hACE2 cells were seeded in 96-well plates (50,000 cells/well) and incubated for approximate 24 hr until reaching over 90% confluent. The pseudovirus were pre-incubated with the supernatant of the circRNA^nAB^ or circRNA^ACE2 decoys^ transfected cells at 37°C for 60 min, and then added to cells in the 96-well plates. Media were changed at 24 hr after transduction. All cells were collected at 48 hr after transduction. Luciferase activity was measured using the Nano-Glo Luciferase Assay System (Promega). The relative luminescence units were normalized to cells infected with supernatant of cell transfected with the circRNA^EGFP^.

### Authentic SARS-CoV-2 NT50 Assay

A549-hACE2 cells were seeded in 96-well plates (20,000 cells/well) and incubated for approximately 24 hr until 90-100% confluent. The mouse serum was 5-fold serially diluted in DMEM, starting at 1:10. The diluted sera were then mixed with titrated virus in a 1:1 (vol/vol) ratio to generate a mixture containing ∼2,000 PFU/well of viruses (MOI = 0.1), followed by an incubation at 37°C for 1 hr. Then the virus/sera mixtures were added to 24-well plates of A549-ACE2 cells, supplemented with 100 µL of DMEM containing 10% FBS in each well. The supernatant and cell pellet precipitation were then collected, and the viral load was detected by RT-qPCR. Briefly, the RNA was extracted from the cell pellet and reverse transcribed. SARS-CoV-2 RNA quantification was performed by RT-qPCR targeting the N gene of SARS-CoV-2 using Roche LightCycler® 96. The abundance of *GAPDH* was set as internal reference. The NT50 was defined as the fold-dilution, which emitted an exceeding 50% inhibition of infection in comparison with the control group.

### Mouse challenge experiments

The mouse model for SARS-CoV-2 Beta variant challenge has previously been characterized^57^. The BALB/c mice immunized with circRNA^RBD-Beta^ (50 µg) were challenged with 5×10^4^ PFU SARS-CoV-2 Beta variant at 7 weeks post boost. The body weight of mice were recorded daily. At 3 days post challenge, the immunized mice were sacrificed, and their lung tissues were collected to measure the viral RNA load, as described below.

### Quantification of viral load in the lung tissues of challenged BALB/c mice by RT-qPCR

The viral RNA load in the lung tissues of challenged mice was detected by quantitative RT-qPCR. Briefly, the lung tissues were collected and homogenized with stainless steel beads in TRIZOL (1 ml for each sample). The RNAs in tissues were then extracted and reverse transcribed. SARS-CoV-2 RNA quantification was performed by RT-qPCR targeting the N gene of SARS-CoV-2, using Roche LightCycler® 96. The abundance of *GAPDH* was set as internal reference. The placebo group viral load was normalized to 100%.

### T cell flow cytometry analysis

The Splenocytes from each immunized mouse were cultured in R10 media (RPMI 1640 supplemented with 1% Pen-Strep antibiotic, 10% HI-FBS), stimulated with RBD peptide pools (**Supplementary Table 1**) at a final concentration of 2 µg/mL for each peptide. 3 hr later, the Golgi Stop transport inhibitor cocktail (BD) was added according to manufacturer instructions. And then, 6 hr later, cells from each group were pooled for stimulation with cell stimulation cocktail (PMA/Ionomycin) as a positive control. Following stimulation, cells were washed with PBS prior to staining with LIVE/DEAD for 20 min at room temperature. Cells were then washed in stain buffer (PBS supplemented with 2.5% FBS) and suspended in Fc Block for 5 min at RT prior to staining with a surface stain of following antibodies: CD3 (Invitrogen, 45-0031-82)/CD4 (BD, 562285)/CD8 (BD, 553035) /CD44 (BD, 563058)/CD62L (BD, 560507). After 20 min, cells were washed with stain buffer, and then fixed and permeabilized using the BD Cytoperm fixation/permeabilization solution kit according to manufacturer instructions. Cells were washed in perm/wash solution, followed by intracellular staining (30 min, RT) using a cocktail of the following antibodies: IFN-γ (BD, 557998)/IL-2 (BD, 560547)/IL-4 (BD, 554435)/TNF-α (BD, 557644). Finally, cells were washed in perm/wash solution and suspended in stain buffer. Samples were washed and acquired on a LSRFortessa (BD Biosciences). Analysis was performed using FlowJo software.

### Ethics and biosafety statement

All rhesus macaques experiments were performed in the animal biosafety level 4 (ABSL-4) facility of National Kunming High-level Biosafety Primate Research Center, Yunnan, China. All animal procedures were approved by the Institutional Animal Care and Use Committee of Institute of Medical Biology, Chinese Academy of Medical Science. Rhesus macaques were monitored at least twice daily throughout the experiment. Commercial monkey chow, treats and fruit were provided daily by trained personnel.

### Rhesus macaque vaccination and plasma collection

For the vaccination of rhesus macaque, groups of 2∼4-year-old rhesus macaques were immunized with LNP-circRNA^RBD^ (20 µg, N= 4; 100 µg, N= 4; 500 µg, N= 4), LNP-circRNA^Ctrl^ (circRNA without the RBD-encoding sequence; 100 µg, N= 4) or PBS (N= 4) in 300 µL (>300 µL in 500 µg dose group) via intramuscular injection in the quadriceps muscle (prime: left, boost: right) twice at a three-week interval. The plasma of immunized rhesus macaques were collected at 0, 1 and 14 days post the prime, and 0, 1, 14, 28 and 35 days post the boost.

### SARS-CoV-2 Virus amplification and identification

Viral stock of native SARS-CoV-2 was obtained from the Center of Diseases Control, Guangdong Province China. Viruses were amplified on Vero-E6 cells and concentrated by ultrafilter system via 300 kD module (Millipore). Amplified SARS-CoV-2 were confirmed via RT-PCR, sequencing and transmission electronic microscopy, and titrated via plaque assay (10^6^ pfu/ml).

### ELISpot assay

The T cell immune responses in rhesus macaques were detected using the PBMCs with commercially available NHP IFN-γ and IL-2 ELISpot assay kits (Mabtech), and NHP IL-4 ELISpot assay kit (U-CyTech). The cryopreserved rhesus macaques PBMCs were thawed and cultured with pre-warmed AIM-V media. For IFN-γ, IL-2 and IL-4 ELISpot assay, 1.0×10^5^ PBMCs were stimulated with final concentration of 1 µg/mL for each RBD peptide (**Supplementary Table 2**). The test for each rhesus macaque were performed in two or three technical repetitions. The dimethyl sulphoxide (DMSO) served as un-stimulated control, and the Phytohemagglutinin (PHA-P, Sigma) and CELL STIMULATION COCKTAIL (ThermoFisher) were used as positive controls. After 24h of stimulation with RBD peptide pools, the streptavidin-HRP substrate (for IFN-γ and IL-2) or AEC substrate (IL-4) were added into plate. The spots were counted by Beijing Dakewei Biotechnology Co., Ltd. The results are background (DMSO treated group) subtracted and normalized to SFC/10^6^ PBMCs.

### SARS-CoV-2 challenge in rhesus macaques

At 5 weeks post boost, all the immunized rhesus macaques were challenged with 1.0×10^6^ PFU of native SARS-CoV-2 virus via the intranasal (0.5 ml) and intratracheal (0.5 ml) routes. The plasma of rhesus macaques was collected and vital clinical signs were recorded on 0, 1, 3, 5 and 7 days post virus challenge. At 7 days post virus challenge, all rhesus macaques were sacrificed to collect specimen for further experiments.

### Histopathology

On the 7 days post virus challenge, the rhesus monkeys were euthanized, and necropsies were performed according to standard protocols. After the dissection, a general examination of the main organs was performed. The lung tissues were harvested, fixed in 10% neutral formalin buffer and embedded in paraffin. 2 µm tissue sections were prepared. Slides were stained with Hematoxylin and Eosin (H&E). The slide images were collected by using Pannoramic DESK and analyzed with Caseviewer C.V 2.3 and Image-Pro Plus 6.0. Histopathological analysis of tissue slides was scored by 3 independent pathologists blinded to the groups of animals.

### Cytokines analyses

The plasma of rhesus monkeys were isolated 24 hr post prime or boost and diluted in 5-fold or 10-fold. All plasma samples were detected using following ELISA kits: IL-6 (Abcam, ab242233), MCP-1 (Cloud-cline corp, SEA087Si96T), TNF-α (Abcam, ab252354), IL-1β (Cloud-cline Corp, SEA563Si96T) and IFN-α (Chenglin, AD0081Mk) according to the manufacturer’s instructions.

### Statistics

An unpaired two-sided Student’s t-test was performed for comparison as indicated in the figure legends. Statistical analyses were performed with Prism 8 (GraphPad Software, Inc.).

### Reporting Summary

Further information on research design is available in the Nature Research Reporting Summary linked to this article.

## Data availability

All data and materials presented in this manuscript are available from the corresponding author (W.W.) upon a reasonable request under a completed Material Transfer Agreement.

